# Divergent roles of orexin and dynorphin in modulating VTA dopamine projections to the BLA after morphine exposure

**DOI:** 10.64898/2026.10.24.761541

**Authors:** Aida Mohammadkhani, Hadi Semizeh, Ijeoma Ifionu, Min Qiao, Stephanie L. Borgland

**Affiliations:** Department of Physiology and Pharmacology, Hotchkiss Brain Institute, The University of Calgary, 3330 Hospital Dr. NW, Calgary, Alberta T2N 4N1 Canada

**Keywords:** opioid dependence, ventral tegmental area, dopamine neurons, orexin, dynorphin, orexin receptor, kappa opioid receptor, basolateral amygdala

## Abstract

**Background:** Opioid use disorder is associated with persistent alterations in mesolimbic dopamine (DA) circuits, but the circuit- and neuromodulatory mechanisms driving this plasticity remain unclear. Lateral hypothalamic orexin/dynorphin (LH_ox/dyn_) neurons provide dense input to the ventral tegmental area (VTA), where orexin and dynorphin exert opposing actions via orexin 1 receptor (Ox1R) and κ-opioid (KOR) receptors, respectively. Here, we investigated how opioid dependence alters the selective contributions of optically driven LH_ox/dyn_ release on the firing of VTA^DA^ neurons that project to the basolateral amygdala (BLA) or the lateral nucleus accumbens shell (lAcbSh).

**Methods:** We measured the effects optogenetic activation of the LH_ox/dyn_ input on VTA^DA^ neuronal activity with retrogradely labeled projection targets after repeated morphine or saline injections or fentanyl self-administration. We used fiber photometry to measure how the LH^ox/dyn^ input to the VTA influences dopamine in the BLA in opioid-dependent mice.

**Results:** We found that in saline-treated mice, LH_ox/dyn_ predominantly inhibited BLA-projecting VTA^DA^ neurons via KOR signaling. Remarkably, opioids, morphine or fentanyl, shifted the neuromodulatory capacity of LH_ox/dyn_ input to BLA-projecting VTA^DA^ neurons from inhibitory to excitatory. This was due to increased dynorphin release in the VTA, leading to reduced function of KORs on BLA-projecting VTA^DA^ neurons. This led to increased BLA^DA^, which typically encodes the emotional and associative salience of reward-related cues in opioid dependent mice.

**Conclusions:** Together, we define a mechanism of how LH_ox/dyn_ corelease independently neuromodulates VTA projections, and how removal of a KOR ‘brake’ leads to enhanced neuromodulatory activity of dopamine projections to the BLA.

## Introduction

The opioid overdose crisis remains a dire public health emergency across North America and surged by 90% during the two years of the COVID-19 pandemic(1). This trend underscores an urgent need for a rigorous mechanistic understanding essential for developing effective interventions that can prevent overdose, reduce mortality, and mitigate long-term harm. Morphine, a prototypic opioid, evokes synaptic plasticity and modifies neuronal activity of dopamine (DA) neurons within the VTA (2). VTA^DA^ neurons integrate diverse excitatory and inhibitory inputs to regulate phasic DA release that encodes reward predictions and drives motivated behavior(3,4). Projection-defined subpopulations of VTA^DA^ neurons mediate distinct aspects of reward processing; neurons projecting to the nucleus accumbens (NAc) primarily drive reinforcement, whereas those projecting to the basolateral amygdala (BLA) encode the emotional and associative salience of reward-related cues(5–7). Because these subpopulations are largely non-overlapping(8), afferents to the VTA can selectively regulate discrete motivational states.

Orexin (hypocretin) neurons in the lateral hypothalamus (LH) provide a major source of modulatory input to the VTA(9–11). LH orexin (ox) fibers densely innervate the VTA and make appositions onto both dopaminergic and GABAergic neurons(9,11). Activation of orexin 1 receptors (Ox1Rs) in the VTA increases intracellular calcium, enhances dopamine neuron firing, and promotes VTA^DA^ release in downstream targets(8,12–14). Functionally, intra-VTA orexin signaling enhances motivation for salient rewards, including palatable food and addictive drugs(15,16). Consistent with this, blockade of orexin receptors reduces both cocaine and opioid self-administration and prevents morphine-induced plasticity in the VTA(17–19). Furthermore, chronic exposure to morphine and fentanyl in rodents leads to a persistent upregulation of orexin peptide expression, with orexin neuron counts elevated in post-mortem hypothalamus from opioid users relative to controls(20–22).

Remarkably, nearly all LH orexin neurons also co-express dynorphin (LH_ox/dyn_), the endogenous ligand for the inhibitory kappa opioid receptor (KOR)(23,24). Orexin and dynorphin are co-packaged within dense-core vesicles and can be co-released(24). Whereas orexin excites VTA^DA^ neurons through Ox1R activation, dynorphin suppresses their activity by activating KORs, which hyperpolarize VTA^DA^ neurons(25,26). While μ opioid receptor (MOR) activation mediates the primary rewarding effects of opioids such as morphine and fentanyl, repeated opioid exposure also leads to upregulation of the dynorphin/KOR system in mesolimbic circuits(27,28).This heightened KOR signaling contributes to negative affective states that arise during opioid withdrawal(28). Thus, LH_ox/dyn_ inputs exert bidirectional effects over VTA^DA^ neurons and their output. Intra-VTA Ox1R antagonism decreases cocaine self-administration, an effect reversed by KOR blockade, suggesting that orexin may oppose the anti-reward actions of dynorphin to facilitate motivated behavior(24). Chronic exposure to opioids such as morphine and fentanyl profoundly alters both the mesolimbic dopaminergic(2,29) and hypothalamic circuits(30,31). Yet, it remains unknown how repeated opioid experience affects the balance between orexin-and dynorphin-mediated signaling in projection-defined VTA subcircuits, and what are the main projection targets of opioid-recruited dopamine populations(32). Here, we tested the hypothesis that opioid exposure differentially influences VTA^DA^ projections modulated by LH_ox/dyn_ input.

## Materials and Methods

### Subjects

All experimental procedures adhered to ethical guidelines established by the Canadian Council for Animal Care and animal use protocols approved by the University of Calgary Animal Care and Use Committee. Adult male and female orexin^cre^ mice (mice expressing the cre recombinase in cells expressing the pre-pro-orexin peptide (post-natal day 60-90) were originally obtained from the Yamanaka lab at the University of Tokyo (1) and bred locally at the University of Calgary Clara Christie Center for Mouse Genomics. Animals used for electrophysiological experiments were group-housed (3-5 per cage) with *ad libitum* access to water and food. Animals used for behavioral experiments were single housed. All Mice were housed in ventilated cages in a temperature (21 ± 2°C) and humidity-controlled (30-40%) room on a 12h reverse light/dark cycle (dark phase from 10:00 AM to 10:00 PM). Electrophysiological experiments were conducted during the animals’ light cycle, whereas all behavioral experiments were performed during the dark cycle.

### Surgical procedures

*Electrophysiology experiments:* All orexin^cre^ mice received bilateral infusions of either channelrhodopsin (‘ChR2’) (AAV2/8-EF1a-DIO-hChR2(H134R)-mCherry; Neurophotonics, Centre de Recherche CERVO, Quebec City, QC, Canada) or control (‘mCherry’) (AAV2/8-hSyn-DIO-mCherry; Neurophotonics) virus. Mice were anaesthetised with 5% isoflurane and with a continuous flow of 2% isoflurane mixed with oxygen thereafter. All measurements were made relative to bregma for viral infusions. Viral injections were performed using a microinjector (Nano-inject II; Drummond Scientific Company, Broomall, PA, USA). Each mouse received 3 bilateral infusions into the LH (100 nl per infusion; 23.1 nl/s; anterior-posterior (AP) −1.35; mediolateral (ML) ± 0.9; dorsoventral (DV) −5.2, −.5,1, −5.0) for a total of 300 nl per hemisphere. After each infusion, the microinjector was left in place for 3 min to allow diffusion of virus away from the needle tip. After all infusions were complete in one hemisphere, the microinjector was left in place for an additional 5 min, 500 mm dorsal of the final injection location to allow diffusion of the virus through the brain tissue. Red retrobeads (max excitation at 530 nm/ max emission at 590 nm) (200 nL; Lumafluor Inc., Naples, FL) were infused bilaterally into the lAcbSh (from bregma: anteroposterior +1.425 mm; mediolateral, ± 1.75 mm; dorsoventral, −4.25 mm) or the BLA (from bregma: anteroposterior −1.0 mm; mediolateral, ± 3.1 mm; dorsoventral, −5.3 mm).

For fiber photometry experiments, AAV9-hsyn-GRAB-DA2m (Titer:2.4×10 GC/ml), a green dopamine biosensor(2), was injected unilaterally into the BLA (from bregma: anteroposterior −1.2 mm; mediolateral, ± 3 mm; dorsoventral, −4.2 mm at 100 nl per infusion; 23.1 nl/s) for a total of 300 nl, while Chrimson, a red-shifted optical actuator(3), (AAV9-Syn-FLEX-ChrimsonR-tdTomato; Titer: 1.8×10 GC/ml) was injected into the LH (100 nl per infusion; 23.1 nl/s; anterior-posterior (AP) −1.35 mm; mediolateral (ML) ±1 mm; dorsoventral (DV) −5.3, −.5,2, −5.1) for a total of 300 nl per hemisphere). Following viral injections, and during the same surgery, orexin^cre^ mice were chronically implanted with an optic fiber (MFC_400/430–0.66_5 mm, Doric Lenses) above the BLA (AP −1.2 mm, ML ±3.0 mm, DV −4.0 mm) to record dopamine, and bilateral optic ferrules (DFC_200/240-0.22, 5.5 mm, Doric lenses) above the VTA (AP −3.5 mm, ML ±0.5 mm, DV −3.8 mm) for optical stimulation of LH_ox/cre_ inputs to the VTA. Implants were then secured to the skull with super glue and dental cement. Injection sites and placements were confirmed in all animals by preparing coronal sections of the lAcbSh and the BLA. All mice received pre- and 3 days postoperative analgesic (meloxicam 5 mg/kg, subcutaneous) and were returned to their home cages and allowed to recover for 6-8 weeks prior to further experimental procedures.

### Morphine Dosing Paradigm

Mice were treated with morphine or saline according to procedures described previously(4). Briefly, repeated intraperitoneal (i.p.) injections of escalating doses of morphine were administered twice daily (AM and PM, 8 h apart) in a procedure known to induce morphine dependence(4). Morphine doses were progressively increased from 10 mg/kg to 50 mg/kg: first day, 2 × 10 mg/kg; second day, 2 × 20 mg/kg; third day, 2 × 30 mg/kg; and fourth day, 2 × 40 mg/kg. On the morning of day 5, mice received a final i.p. injection of 50 mg/kg morphine. Control mice were treated with 0.9% sterile saline following the same procedure. All electrophysiological experiments were performed 2 h following the last morphine or saline injection.

### Electrophysiology

All electrophysiological recordings were performed in slice preparations from adult male and female orexin^cre^ mice 6-8 weeks after receiving the cre-dependent viral vector containing ChR2. Briefly, mice were anaesthetized with isoflurane and transcardially perfused with an ice-cold N-methyl-d-glucamine (NMDG) solution containing (in mm): 93 NMDG, 2.5 KCl, 1.2 NaH2PO4.2H2O, 30 NaHCO3, 20 HEPES, 25 D-glucose, 5 sodium ascorbate, 3 sodium pyruvate, 2 thiourea, 10 MgSO4.7H2O and 0.5 CaCl2.2H2O and saturated with 95% O2–5% CO2. Mice were then decapitated, and brains were extracted. Sections containing the retrobead injection site were confirmed visually. Horizontal sections (250 μm) containing the VTA were cut in NMDG solution using a vibratome (VT1200, Leica Microsystems, Nussloch, Germany). Slices were then incubated in NMDG solution (32°C) saturated with 95% O2–5% CO2 for 10 min before being transferred to a holding chamber containing artificial cerebrospinal fluid (ACSF), (in mM): 126 NaCl, 1.6 KCl, 1.1 NaH2PO4, 1.4 MgCl2, 2.4 CaCl2, 26 NaHCO3 and 11 glucose (32–34°C) equilibrated with 95% O2–5% CO2 for at least 45 min before recording. Slices were transferred to a recording chamber on an upright microscope (Olympus BX51WI) and continuously superfused with ACSF (2 mL/min, 34 °C). Cells were visualized on an upright microscope using “Dodt-type” gradient contrast infrared optics and whole-cell recordings were made using a MultiClamp 700B amplifier (Axon Instruments, Molecular Devices). Recording electrodes (3-5 MΩ) for measuring firing rates were filled with potassium-D-gluconate internal solution (in mM): 130 potassium-D-gluconate, 4 MgCl2, 10 HEPES, 0.5 EGTA, 10 sodium creatine phosphate, 3.4 Mg-ATP and 0.3 Na2GTP and 0.2% biocytin.

Putative dopamine neurons were identified by their location, fusiform shape and the presence of a hyperpolarization-activated cation currents (*I*_h_). In some experiments, dopamine neurons were identified by their presence of retrograde label and posthoc co-labeling for tyrosine hydroxylase (TH; see below). After breaking into the cell, I_h_ was recorded in voltage-clamp mode using a voltage step to −130 mV to dopamine neurons voltage-clamped at −70 mV. I_h_ was determined as the change in current between ∼30 ms and 248 ms after the voltage step was applied. Because most dopamine neurons ceased firing within 5 min of recording, current-step induced firing was used for all experiments. For current-step experiments, the membrane potential for each neuron was set to −60 mV by DC injection via the patch amplifier and a series of 5 current pulses (250 ms in duration, 5-25 pA apart, adjusted for each cell) were applied every 45 s, where the minimum current amplitude was set for each cell so that the first pulse was subthreshold and did not yield firing. From this series of current steps, we then selected a current step that yielded 3-5 action potentials during the baseline period and used that step for the analysis of peptide effects, as described previously(5,6).

All recordings were performed in synaptic blockers to assess the role of peptide neuromodulation rather than that of fast transmitters. The recording solution included AP5 (50 μM), DNQX (10 μM), and picrotoxin (100 μM), antagonists of NMDA receptors, AMPA receptors, and GABA_A_ receptors, respectively. After obtaining a stable baseline of firing for 10 min, we optically stimulated LH_ox/dyn_ inputs in the VTA once at 30 Hz, 30 s (5 ms pulse width, 5 mW), a frequency and duration previously shown to produce the largest change in firing(6), from a blue light-emitting diode (LED) (470nm) directly delivered through the light path of the Olympus 40X water immersion lens. 5 min after stimulation, either SB334867 (Ox1R antagonist, 1 µM, Tocris) or NorBNI (KOR antagonist; Tocris; 1 µM) were applied for 15 min. In morphine-treated mice, acute brain slices were prepared 2 h after the last injection of morphine and maintained in vitro in morphine sulphate (5 μM) throughout the experiments (in holding chamber and throughout recordings) to prevent spontaneous opioid withdrawal(7,8). In the other experiments, slices from morphine-treated animals were maintained in normal morphine-free ACSF after slicing (for 1–6 h). We recorded the response of VTA^DA^ neurons, identified by electrical characteristics and posthoc TH staining, after optical stimulation of LH_ox/dyn_ inputs in the VTA, using whole cell patch clamp electrophysiological recordings. Recordings were performed from three experimental groups: slices from saline-treated mice, slices from morphine-treated mice maintained in morphine-free ACSF, and slices from morphine-treated mice maintained in ACSF containing 5 μM morphine. Exogenous morphine was added to the bath and recording solutions to prevent spontaneous withdrawal, which can contribute to increased neuronal activity (9).

For G-protein inwardly rectifying potassium channel (GIRK) and holding potential recordings, tetrodotoxin (TTX; 500 nM) was added to block neural activity and U69593 (5 µM) subsequently added to this TTX solution. In GIRK experiments, current–voltage data were collected in voltage clamp by stepping from a holding potential of −60 to −40 mV and ramping down to −140 mV over a 2 s interval(10).

### Enzyme-Linked Immunosorbent Assay (ELISA) for peptides

To examine dynorphin or orexin release in the VTA, brain slices from LH_ox/dyn_ ChR2-expressing morphine or control mice were prepared as described above. Based off a procedure measuring electrically evoked orexin and dynorphin release with ELISAs (11), a mini-slice of the VTA containing LH_ox/dyn_ terminals was dissected from the midbrain slice and placed into a small culture dish with 200 mL of ACSF. A laser-connected fibre optic (Doric Lenses; 200 µm) was placed directly above the VTA mini-slice, and ChR2-containing LH_ox/dyn_ terminals were optically stimulated once at 30 Hz, 30 s (5 mW power) to evoke neuropeptide release. Supernatant from slices (200 µl) was collected 5 min after optical stimulation, snap frozen, and stored at −80 C. Samples were then tested with an ELISA assay for either dynorphin-A (1-17; Cat# EKU02719-96T) or orexin-A (Cat#EKC37520-96T) (Biomatik, Canada) in accordance with the manufacturer’s instructions.

### RNAscope Multiplex Fluorescent Assay

RNAScope was performed using the RNAscope Multiplex Fluorescent Reagent Kit v2 and probes from Advanced Cell Diagnostics (ACD) as per manufacturer’s instructions. Briefly, frozen brains were sectioned to 10-20 μm using a cryostat and mounted onto Superfrost Plus slides (Fisher Scientific). Sections were fixed with cold 4 % paraformaldehyde for 1 h at 4 °C and then dehydrated in increasing concentrations of ethanol (50, 70, and 100 % twice) for 5 min at each concentration. Sections were treated with hydrogen peroxide for 10 min, followed by RNAScope protease IV incubation for 30 min at room temperature. Probes were hybridized and signals amplified as indicated in the manufacturer’s instructions. The mouse *Hcrt1* probe (Cat#490461-C2) (ACD, USA) was assigned to channel 2 (C2), and the mouse kappa *Oprk1* probe (Cat# 316111-C3) (ACD, USA) was assigned to channel 3 (C3). The C2 and C3 probes were diluted 1:50 with the recommended probe diluent. Signal development was performed sequentially for each channel with Opal 520 and 570 fluorophores (diluted 1:1500) to visualize probes. DAPI was used for counterstaining nuclei. Coverslips were applied with Fluoroshield mounting medium (MilliporeSigma, Canada). Imaging was performed using an Olympus Virtual Slide Scanner with a 10× objective and a Leica SP8 confocal microscope with a 40x objective. Quantification of positive *Hcrt1*, *Oprk1*, and colocalized-positive cells was conducted using QuPath software. Cell counts included the total number of retrobead labeled VTA neurons expressing *Hcrt1* and/or *Oprk1* mRNA. Copies of mRNA within each cell (puncta) and neurons with positive mRNA expression were analyzed to assess expression levels and the extent of colocalization.

### In Vivo fiber photometry recording and analysis

Fiber photometry was performed using a Doric Lenses dual-channel photometry console coupled to a dual LED driver. Two excitation wavelengths were delivered: 465 nm (for GRAB_DA_2m excitation) and 405 nm (isosbestic control). Fluorescence emission (500–550 nm) was collected through a dichroic Mini Cube and directed to photodetectors (Newport Visible Femtowatt Photoreceiver Module Model 2151). Light power was calibrated to deliver 30 μW light at the tip of the fiber using a photometer (Thor Labs). Raw fluorescence signals were demodulated in real time (cut off frequency for demodulated signal: 12 Hz). Using custom MATLAB analyses, the 405 nm channel was used as an isosbestic control, and the data recorded using this channel was fit to a biexponential decay. Data from the Ca^2+^-dependent 465 nm channel were then linearly scaled using the fitted decay of the isosbestic channel, correcting for any photobleaching which occurred during the recordings(12). The resulting vector was converted to a ΔF/F trace, and each trace was normalized as a z-score. Trials containing abrupt movement-related artifacts exceeding 3 standard deviations from baseline were excluded from analysis. ΔF/F traces were down-sampled to 10 Hz for further analysis. For z-score, the first 600 s of baseline were used to estimate mean and SD. Z-scored ΔF/F traces were analyzed for area under the curve, peak frequency, and maximum fluorescence. Area under the curve (AUC) was calculated within predefined stimulation epochs corresponding to the 30 s laser trains.

Mice were treated with repeated saline or morphine as described above. Two hours after the final morphine or saline injection, bilateral optic ferrules were coupled to diode-pumped solid-state lasers (589 nm for crimson opsin activation; 473 nm for control). Light trains (30 Hz, 5 ms pulse width, 30 s duration) were generated by a Pulse Pal stimulator (Sanworks) synchronized with the Doric system and delivered every 5 min for a total of three trains per 15-min session. Laser output was calibrated to 10–12 mW at the ferrule tip. Laser output was monitored before and after each recording session to ensure stable power delivery across animals. Data were exported from the Doric system and processed in MATLAB. 405 nm signals were used to correct 465 nm signals via linear regression, producing ΔF/F traces. Z-score normalization was applied using baseline (0–600 s). In some experiments, the day after the first photometry recording, mice received a 10 min baseline photometry recording, an injection of morphine (50 mg/kg, i.p.), a 30 min photometry recording, and then an injection of naloxone (5 mg/kg, i.p.) and an additional 10 min recording.

### Oral fentanyl self-administration

Self-administration training occurred in Med Associates operant chambers (Med Associates, VT, USA), fitted with 2 levers (active and inactive), a stimulus light located directly above the active lever, and house light. Operant boxes were housed in individual sound-attenuating chambers, and all boxes were controlled by Med-PC IV software (Med Associates). Self-administration training was performed in 2 h sessions on a fixed-ratio (FR-1) schedule. Mice were habituated to the room and then given magazine training for 1 day. Mice were then trained on an FR-1 schedule for an oral solution of sucrose whereby active lever presses delivered 70 µl of 10%, sucrose, paired with a cue light above the lever, and followed by a 10 s time-out signaled by the termination of the cue light. In sucrose self-administering mice, mice continued daily FR-1 sessions 5 days per week for 15 sessions. For oral fentanyl self-administering mice, after mice were trained to a criterion of > 25 reinforced presses for 3 days on sucrose, mice were then switched to an oral fentanyl solution. A response on the active lever resulted in a 2 s infusion of 10 µg/ml (70 µl) of fentanyl citrate solution to the magazine, paired with light above the lever. Each liquid delivery was followed by a 10 s time-out signaled by termination of the cue light. Responses on the inactive lever had no consequences. Training sessions terminated after 2 h. Mice were trained once daily 5 days/week for 20 self-administration sessions and to a criterion of >25 infusions for three consecutive sessions. Two hours after the last fentanyl or sucrose self-administration session, mice were taken for brain slice preparation (described above) or naloxone precipitated withdrawal (described below).

### Behavioral Assessment of Naloxone Precipitated Withdrawal

Naloxone (2 mg/kg, naloxone hydrochloride dihydrate, Sigma) was delivered 2 h after the final oral fentanyl or sucrose self-administration session to precipitate opioid withdrawal. Naloxone-or saline-precipitated withdrawal signs were scored over the course of 30 min (in 5 min intervals), as previously described(13). Briefly, jumping, headshakes, ‘wet-dog’ (full-body) shakes, chewing/licking, and teeth chattering were given a composite score of 0–3 for each 5 min interval (0 = absent, 1 = 1-3 bouts; 2=4-6 bouts, 3 = 7 bouts and greater). See supplemental Table 1. The presence of piloerection, salivation, and paw tremors were also scored every 5 min and assigned one point. To calculate weight loss, mice were weighed before and after the 30 min scoring period as an indicator of micturition and defecation. All behavioral and physiological signs were summed to generate a cumulative withdrawal score(13).

### Immunohistochemistry and confocal microscopy

To identify dopamine neurons, we filled neurons with biocytin electrophysiological whole-cell recordings and subsequently processed slices for tyrosine hydroxylase (TH)- like immunoreactivity, as described previously(6). Brain slices from patch-clamp recordings in orexin^cre^ mice were fixed overnight in cold 4% paraformaldehyde and then stored in phosphate-buffered saline until processing. Sections were then blocked in 10% normal goat serum and incubated with mouse anti-TH (1:1000) for 24 hours at room temperature. Alexa Fluor 488 goat anti-mouse (1:400) (Life Technology, A21236) and Dylight 594 conjugated streptavidin (Jackson Immuno,016-580-084) were applied to identify dopamine neurons tagged with biocytin. Slices were mounted with Fluoroshield.

To check for co-localization of ChR2 expression and orexin in the LH, mice were deeply anesthetized with isoflurane and transcardially perfused with phosphate buffered saline (PBS) and then with 4% paraformaldehyde (PFA). Brains were dissected and post-fixed in 4% PFA at 4°C overnight, then switched to 30% sucrose. Coronal frozen sections were cut at 30 µm using a cryostat. 10% goat serum was applied to block non-specific binding for 1 h. Sections were then incubated with primary antibody rabbit anti-orexin 1:500 (Phoenix pharmaceuticals, H-003-30) and chicken red fluorescent protein (RFP) 1:2000 (Rockland, 600-901-379) in 1% BSA at 4°C overnight followed by incubation with secondary antibody Alexa Fluor 488 goat anti-rabbit (Life Technology, A-21206) and Alexa Fluor 594 goat anti chicken (Life Technology, A-11042) 1:400 for 1 h.

To assess for the colocalization of retrobeads (Lumifluor), biocytin and TH in VTA patch-clamped slices, 10% goat serum was applied to block non-specific binding for 1 h. Sections were then incubated with primary antibody mouse anti-TH (1:1000) in 1% BSA for 24 h at room temperature followed by incubation with secondary antibody Alexa Fluor 647 goat anti mouse (1:400) and Alexa fluor 488 streptavidin (1:200) for 2 h.

All images were obtained at 10x on an Olympus scanner microscope (Olympus Canada, Ontario, Canada) and at 20x or 40x on a Leica SP8 confocal microscope (Leica Biosystem Canada Inc. Ontario, Canada). Quantification of orexin expressing cells in the LH was performed with QuPath.

### Drugs

Morphine sulphate (Toronto Research Chemicals, Canada) and naloxone HCl (Sigma, USA) were dissolved in physiological saline (0.9%). SB334867 (1 µM), Nor- binalthorphin (NorBNI, 1 µM), or U69594 (5 µM) (Cayman Chemical, USA) were dissolved in DMSO and then diluted to their final concentration containing 0.001% DMSO in ACSF and TTX citrate (0.5 µM, Hello bio, USA) was dissolved in ACSF and bath applied at a rate of 2 ml/min to midbrain slices for electrophysiological experiments. The short-acting, reversible KOR antagonist, aticaprant (LY2456302) (Cayman Chemical, USA) was diluted in a vehicle of 10% DMSO and 90% (20% (w/v) sulfobutylether-β-cyclodextrin (SBE-β-CD) in saline). Aticaprant (10 mg/kg, i.p.) was administered twice a day for 5 days in mice exposed to morphine or saline.

### Data analysis and statistics

All data are presented at mean ± SEM, and each data point represents an individual response from a neuron or animal. Firing data for all neurons was analyzed with the MiniAnalysis program (Synaptosoft). Optical stimulation-induced changes in firing are expressed as a percentage of the averaged 10 min baseline. Additionally, a comparison of the number of action potentials (APs) per a current step were averaged during the last 2 min of baseline, a 2 min period during the 5 min period after stimulation, and a 2 min period at the end of the recording was performed using a one-way ANOVA. To distinguish responders showing a decrease or increase from non-responders, we used a criterion of a 20% change in firing rate from the baseline. Responses from neurons of male and female mice were analyzed together. Our pilot data indicated that there were not sufficiently large sex differences in electrophysiological responses and therefore mice were grouped together as these studies were not sufficiently powered to test for sex differences. All statistical analyses were performed in GraphPad Prism 9.4.1 (GraphPad, US). Paired t-tests were used to compare before and after drug applications whereas a student t-test was used to compare between two independent groups (eg peptide responses). In the case where 3 or more timepoints were compared, a repeated measures one-way ANOVA was used with a Dunnett multiple comparisons test. Alternatively, if groups were independent a one-ANOVA with Holm–Šídák multiple comparison test was used. To compare if there was a change in proportion of the number of cells recorded increasing vs decreasing their responses, we used a Fisher’s Exact test. In all electrophysiology experiments, sample size is expressed as n/N where “n” refers to the number of cells recorded from “N” animals. For visualization and statistical analysis of photometry experiments, ΔF/F and z-score data were imported from MatLab (Mathworks,) into GraphPad Prism. Area under the curve (AUC), peak amplitude, and time-to-baseline were quantified. Group data are presented as mean ± SEM. Sample sizes are consistent with those reported in similar studies and provide sufficient power to detect changes with the appropriate statistical analysis. Data used for statistical comparison fit a Gaussian distribution, and variances were equivalent between groups, unless otherwise stated. All significance was set at P<0.05. The levels of significance are indicated as follows: ****p<0.0001, ***p< 0.001, **p < 0.01, *p < 0.05. Descriptive statistics are presented in the figure legends.

## Results

### More VTA^DA^ neurons increase firing to optical stimulation of LH_ox/dyn_ inputs in morphine-dependent mice

Opioids influence LH_ox/dyn_ neurons by either suppressing firing(33,34) or increasing orexin peptide expression and thus turn on new orexin neurons with chronic opioid use(20,21). Thus, we asked how opioid dependence impacts how LH_ox/dyn_ inputs influence VTA^DA^ neuron firing. Mice received twice-daily escalating doses of morphine (Figure S1A), which robustly produces naloxone precipitated opioid withdrawal (35,36) (Figure S1B-I). To study the effect of opioid dependence on LH_ox/dyn_ regulation of VTA^DA^ neuronal activity, we expressed ChR2 in LH_ox/dyn_ neurons of orexin^cre^ mice and photoactivated fibers within the VTA (Figure 1A-C). Because dynorphin is expressed in orexin-containing neurons and no other LH neurons (23,24), this manipulation is specific to the LH_ox/dyn_ population and both peptides are released (14,37). To eliminate potential effects of co-released glutamatergic transmission (38), evoked firing of VTA^DA^ neurons was recorded in the presence of synaptic blockers. Following a baseline, we optically stimuled LH_ox/dyn_ inputs to the VTA (at a 30 Hz, 30 s), which increased or decreased the firing of VTA^DA^ neurons, identified by post-hoc biocytin staining (Figure 1D) (14,39). If an increase in firing frequency occurred in response to optical stimulation, we applied the Ox1R antagonist, SB334867 (1 μM, 15 min), as orexin is known to increase firing of VTA^DA^ neurons(8,12,14). Conversely, if a decrease in firing occurred, we administered the KOR antagonist, NorBNI (1 μM, 15 min), as dynorphin is known to decrease firing of VTA^DA^ neurons(8,25). We did not apply an antagonist if no change in firing occurred (Figure 1F).

**Figure 1.**
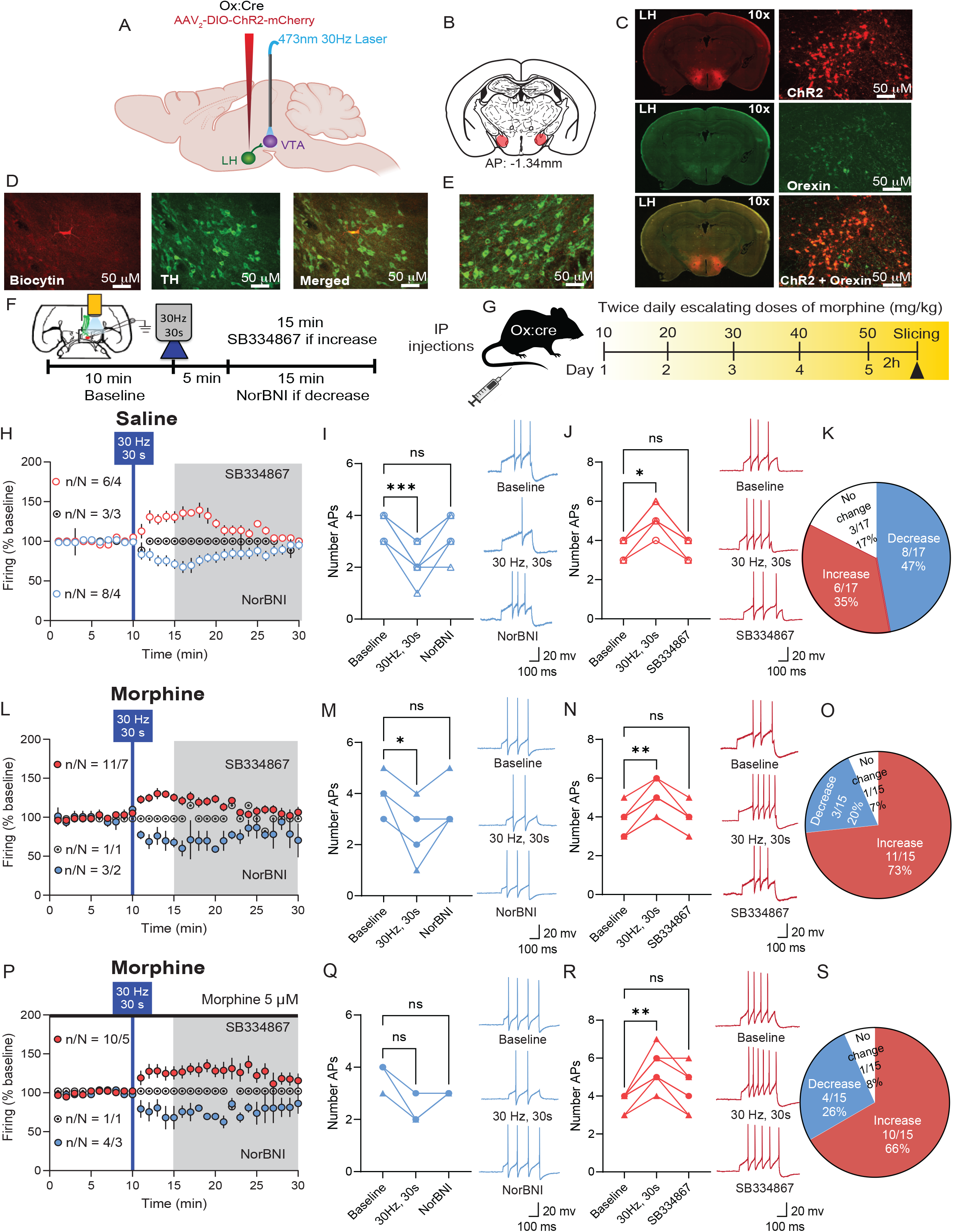
Firing of dopamine neurons in response to optical stimulation of LH_ox/dyn_ inputs increases in morphine-dependent mice. (A) Schematic of viral strategy. LH, lateral hypothalamus; VTA: ventral tegmental area. (B) Coronal brain sections mapping of ChR2 expression areas within the LH (−1.34mm AP from Bregma). (C) Immunohistochemical demonstration of cre-dependent expression of ChR2-mCherry (red, upper), orexin (green, middle), and a merged image of orexin and mCherry (ChR2 reporter) within the LH (lower). (D) Identification of VTA^DA^ neurons co-labeled with biocytin (red, left) and tyrosine hydroxylase (TH, green, middle); merged image shows a biocytin-filled TH-positive neuron (right). (E) Example image of LH_ox_ fibers expressing ChR2 within the VTA of orexin^cre^ mice (merged image of VTA^DA^ neuron labeled with TH (green) and LH_ox_ fibers expressing ChR2 (mCherry reporter, red). (F) Experimental timeline of electrophysiology experiments with optical stimulation. (G) Timeline of twice-daily escalating morphine treatments. (H) Time course of evoked firing before and after 30 Hz, 30s photoactivation of LH_ox/dyn_ inputs in VTA^DA^ neurons of ChR2-expressing saline-treated mice (total recorded neurons: n/N=17/11). Optical stimulation of LH_ox/dyn_ inputs in the VTA produce a diverse response on firing rate of VTA^DA^ neurons in saline-treated mice. (I-J) Number of action potentials (APs, left) and sample traces (right) recorded during baseline, after 30 Hz, 30 s optical stimulation, and following NorBNI or SB334867 application, decreased (I; n/N=8/4, Repeated measures (RM) one-way ANOVA: F_(2, 12)_=18, p=0.010) or increased (J; n/N=6/4, F_(1, 5)_= 45, p=0.019) responses to LH_ox/dyn_ optical stimulation in VTA^DA^ neurons. Each symbol and line represent an individual neuron recorded during the baseline, after 30Hz, 30s stimulation and in the presence of norBNI (I) or SB334867 (J). Some symbols may be occluded. (K) Proportion of increased or decreased responses of VTA^DA^ neurons to LH_ox/dyn_ photostimulation. (L) Time course of evoked firing showing decreased or increased responses to LH_ox/dyn_ optical stimulation of morphine-treated mice (total number of cells recorded: n/N=15/10). (M,N) Number of action potentials (left) and sample traces (right) recorded during baseline, after optical stimulation, and following NorBNI or SB334867 application, decreased (M; n/N=3/2, RM one-way ANOVA: F_(2, 6)_ =7, p=0.037) or increased (N; n/N=11/7, RM one-way ANOVA: F _(1, 12)_=49, p=0.0036) responses to LH_ox/dyn_ optical stimulation in VTA^DA^ neurons of morphine-treated mice. Each symbol and line represent an individual neuron recorded during the baseline, after 30Hz, 30s stimulation and in the presence of norBNI (M) or SB334867 (N). Some symbols may be occluded. (O) There was a shift to a greater proportion increased responses of VTA^DA^ neurons to LH_ox/dyn_ photostimulation in morphine-compared to saline-treated mice (χ²=5.04, p=0.025). (P) Time course of evoked firing before and after photoactivation of LH_ox/dyn_ inputs in VTA^DA^ neurons of morphine-treated mice in the presence of morphine in the recording solution (total number of cells recorded: n/N=15/9). (Q, R) Number of action potentials and sample traces that showed decreased (Q; n/N=4/3, F_(2, 4)_ =8 p=0.083) or increased (R; n/N=10/5, RM one-way ANOVA: F_(2, 11)_ = 19.20, p=0.0042) responses to LH_ox/dyn_ optical stimulation. Each symbol and line represent an individual neuron recorded during the baseline, after 30Hz, 30s stimulation and in the presence of norBNI (Q) or SB334867 (R). Some symbols may be occluded. (S) There was a greater proportion increased responses of VTA^DA^ neurons to LH_ox/dyn_ photostimulation in morphine-treated (with morphine in recording solution) compared to saline-treated mice (χ²=4.47, p=0.034). Data presented as mean ± SEM or individual values from male (circles) or females (triangles). ∗p < 0.05, ∗p < 0.05. Scale bars, 50 μm.

Optical stimulation of LH_ox/dyn_ inputs to VTA^DA^ neurons of saline-treated mice significantly decreased firing in 47% of VTA^DA^ neurons (Figure 1H-K), an effect mediated by KORs (Figure 1I) and significantly increased firing in 35% of VTA^DA^ neurons (Figure 1J-K), an effect mediated by Ox1R (Figure 1J). The proportion of cells activated by optical stimulation of LH_ox/dyn_ fibers significantly increased in morphine- treated mice compared to saline treated mice (Figure 1K,O, χ²=5.04, p=0.025). Seventy-three percent of VTA^DA^ neurons had significantly increased firing and 20% were significantly decreased by photoactivation of LH_ox/dyn_ fibers in the VTA (Figure 1L-O), suggesting that repeated morphine exposure shifts the balance of orexin/dynorphin signaling toward excitation. We assessed if this effect was due to an excitatory effect of opioid withdrawal occurring in the slice(40) by maintaining neurons in morphine throughout the ex vivo environment(40,41). Notably, photoactivation of LH_ox/dyn_ fibers predominantly increased firing in the presence of continued morphine compared to saline treated mice (Figure 1P-S, χ²=4.47, p=0.034). Together, these results indicate that repeated morphine exposure enhances LH_ox/dyn_ excitatory control over VTA^DA^ neurons, both in the absence and continued presence of morphine.

### More BLA-projecting VTA^DA^ neurons increase firing to optical stimulation of LH_ox/dyn_ inputs in morphine-dependent mice

LH_ox/dyn_ differently activates VTA^DA^ neurons depending on the projection target. VTA^DA^ neurons projecting to the lAcbSh predominantly increase firing to LH_ox/dyn_ stimulation, an effect driven by orexin. In contrast, VTA^DA^ neurons projecting to the BLA decrease firing to LH_ox/dyn_ stimulation, an effect driven by dynorphin(42). To determine whether repeated morphine differentially alters LH_ox/dyn_ modulation of specific VTA^DA^ subpopulations, we recorded lAcbSh-projecting VTA^DA^ identified by retrograde labeling (Figure 2A–C). In saline-treated mice, optical stimulation of LH_ox/dyn_ terminals increased firing in most lAcbSh-projecting VTA^DA^ neurons, consistent with previous findings of orexin-mediated excitation in naïve mice(14). Inhibitory (Figure 2E) and excitatory (Figure 2F) responses were washed off by NorBNI or SB334867, confirming dynorphin or orexin contributions, respectively. In morphine-treated mice, regardless of whether slices were maintained in morphine-free ACSF (χ²=0.21, p=0.65; Figure S2A-E) or in ACSF containing morphine (χ²=0.92, p=0.34; Figure 2H-K), there was no difference in proportion of lAcbSh-projecting VTA^DA^ neurons activated or inhibited by LH_ox/dyn_ stimulation compared to saline-treated mice. These findings indicate that repeated morphine exposure does not alter LH_ox/dyn_ modulation of lAcbSh-projecting VTA^DA^ neurons.

**Figure 2.**
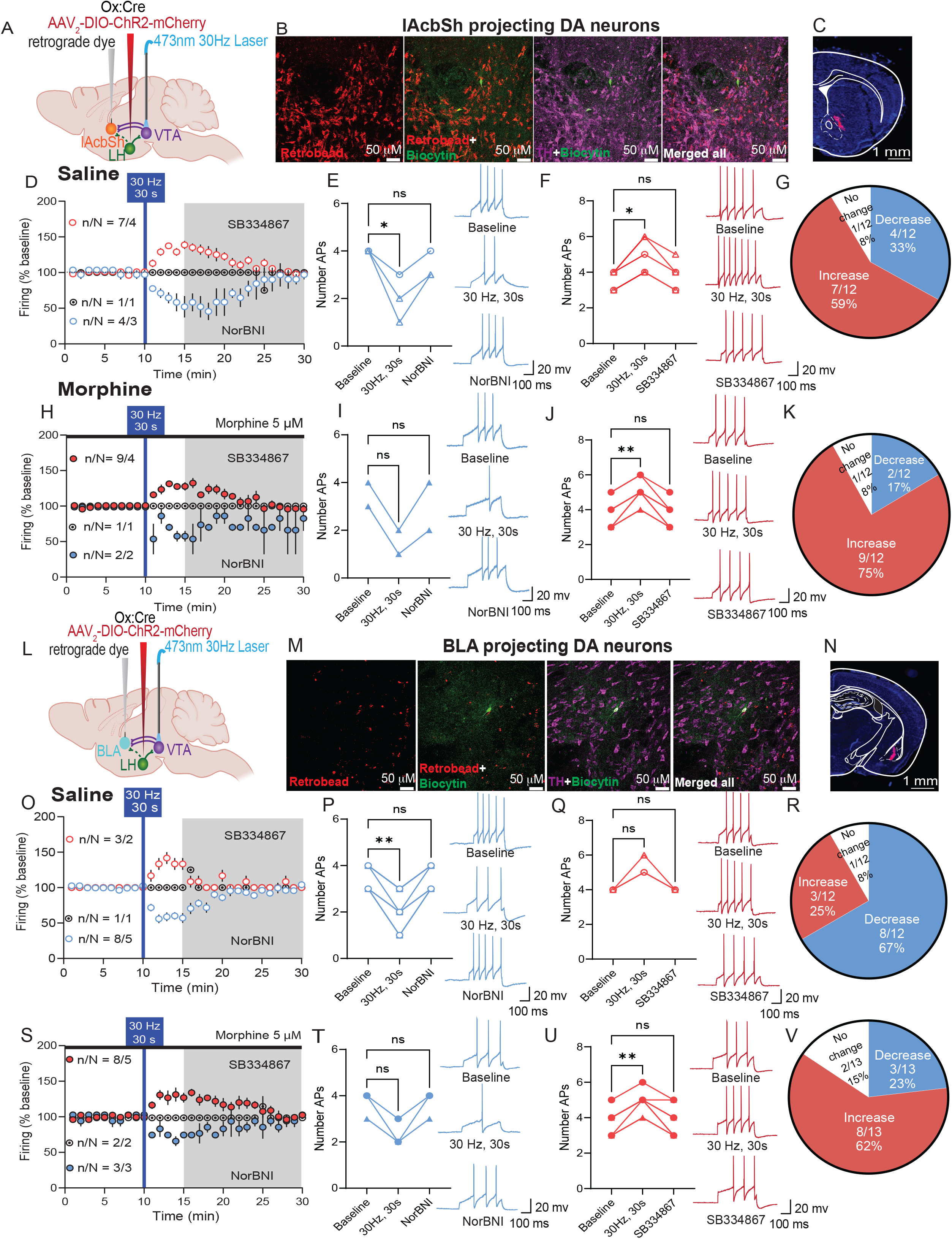
Repeated morphine increases firing of BLA-projecting dopamine neurons in response to optical stimulation of LH_ox/dyn_ inputs, but does not change the neuromodulatory capacity of LH_ox/dyn_ on lAcbSh-projecting VTA^DA^ neurons. (A) Schematic of the viral strategy. (B) Images of VTA sections with TH positive neurons (purple), biocytin (green), and retrograde labeled (red) neurons from the lAcbSh. (C) Example of retrobead injection site in the lAcbSh of orexin^cre^ mice. (D) Time course of evoked firing before and after photoactivation of LH_ox/dyn_ inputs onto lAcbSh-projecting VTA^DA^ neurons of saline-treated mice (total number of cells recorded: n/N=12/8). (E-F) Number of action potentials and sample traces during baseline, after optical stimulation, and after NorBNI or SB334867 (1 µM) application in lAcbSh-projecting VTA^DA^ neurons that showed decreased (F; n/N=4/3, RM one-way ANOVA: F_(2, 4)_=11.20, p=0.049) or increased (G; n/N=7/4, RM one-way ANOVA: F_(2,8)_=24.57 p=0.023) responses to LH_ox/dyn_ optical stimulation. Each symbol and line represent an individual neuron recorded during the baseline, after 30Hz, 30s stimulation and in the presence of norBNI (E) or SB334867 (F). Some symbols may be occluded. (G) Optical stimulation of LH_ox/dyn_ inputs in the lAcbSh-projecting VTA^DA^ preferentially increased firing rate of VTA^DA^ neurons in saline-treated mice. (H) Time course of evoked firing before and after photoactivation of LH_ox/dyn_ inputs onto lAcbSh-projecting VTA^DA^ neurons of morphine-treated mice with morphine in the recording solution (total number of cells recorded: n/N=12/7). (I, J) Number of action potentials and sample traces that showed few neurons decreased (N; n/N=2/2, RM one-way ANOVA: F_(2, 2)_=13, p=0.16), but most increased (O; n/N=9/4, RM one-way ANOVA: F_(2, 16)_=53, p=0.0044) responses to LH_ox/dyn_ optical stimulation in the presence of morphine in the recording solution. Each symbol and line represent an individual neuron recorded during the baseline, after 30Hz, 30s stimulation and in the presence of norBNI (I) or SB334867 (J). Some symbols may be occluded. (K) In morphine-treated mice, while photoactivation of LH_ox/dyn_ inputs primarily increases firing rate of lAcbSh-projecting VTA^DA^ neurons maintained in morphine, it is not different from saline-treated mice (χ²=0.92, p=0.34). (L) Schematic of viral strategy. (M) Images of VTA sections with TH positive neurons (purple), biocytin (green), and retrograde labeled (red) neurons from the BLA. (N) Example of a retrobead injection site in the BLA of orexin^cre^ mice. (O) Time course of evoked firing before and after photoactivation of LH_ox/dyn_ inputs in BLA-projecting VTA^DA^ neurons of saline-treated mice (total number of cells recorded: n/N=12/8). (P,Q) Number of action potentials and sample traces during baseline, after optical stimulation, and after NorBNI or SB334867 application in BLA-projecting VTA^DA^ neurons that showed decreased (P; n/N=8/5, RM one-way ANOVA: F_(2, 14)_ = 56.47, p=0.0054) or increased (Q; n/N=3/2, RM one-way ANOVA: F_(2, 4)_= 16, p=0.13) responses to LH_ox/dyn_ optical stimulation. Each symbol and line represent an individual neuron recorded during the baseline, after 30Hz, 30s stimulation and in the presence of norBNI (P) or SB334867 (Q). Some symbols may be occluded. (R) Optical stimulation of LH_ox/dyn_ inputs in the BLA-projecting VTA^DA^ preferentially decreased firing rate of VTA^DA^ neurons in saline-treated mice. (S) Time course of evoked firing before and after photoactivation of LH_ox/dyn_ inputs in BLA-projecting VTA^DA^ neurons of morphine-treated mice in the presence of morphine in the recording solution (total number of cells recorded: n/N=13/10). (T, U) Number of action potentials and sample traces that showed fewer decreased (T; n/N=3/3, RM one-way ANOVA: F_(2, 4)_=16, p=0.13) and more increased (U; n/N=8/5, RM one-way ANOVA: F_(2, 14)_= 24.65, p=0.0081) firing responses to photoactivation of LH_ox/dyn_ inputs to BLA-projecting VTA^DA^ neurons, in the presence of continued morphine. Each symbol and line represent an individual neuron recorded during the baseline, after 30Hz, 30s stimulation and in the presence of norBNI (T) or SB334867 (U). Some symbols may be occluded. (V) In morphine-treated mice, photoactivation of LH_ox/dyn_ inputs primarily increases firing rate of BLA-projecting VTA^DA^ neurons in the presence of continued morphine compared to saline-treated mice (χ²=4.55, p=0.033). Data presented as mean ± SEM or individual values from male (circles) or females (triangles). ∗p < 0.05, ∗∗p < 0.01. Scale bars, 50 μm (B, M). Scale bars, 1 mm for (C, N).

We next examined how LH_ox/dyn_ optical stimulation influenced BLA-projecting VTA^DA^ neurons from repeated morphine or saline exposed mice (Figure 2L-N). In saline-treated mice, optical stimulation of LH_ox/dyn_ inputs to BLA-projecting VTA^DA^ neurons predominantly decreased firing (Figure 2O-R), an effect washed off by norBNI, consistent with dynorphin-mediated inhibition. Remarkably, in morphine-dependent mice, photoactivation of LH_ox/dyn_ inputs to BLA-projecting VTA^DA^ neurons predominantly increased firing both in the absence (Figure S2F–H) or presence of morphine in the recording solution (Figure 2S-V). This activity shift in morphine-dependent mice was significantly different from saline-treated mice (ACSF: χ²=5.24, p=0.022, morphine in ACSF: χ²=4.55, p=0.033). Thus, repeated morphine exposure changes the response of BLA-projecting VTA^DA^ neurons to LH_ox/dyn_ stimulation from inhibition to excitation. This shift suggests that repeated opioid exposure may enhance orexin influence and/or reduce dynorphin signaling onto BLA-projecting VTA^DA^ neurons.

Morphine increases excitability of VTA^DA^ neurons (43,44), therefore we assessed if there were differences in baseline excitability (pre optical stimulation of LH_ox/dyn_ inputs) between lAcbSh-or BLA-projecting VTA^DA^ neurons from saline or morphine treated mice. Notably, overall lAcbSh-projecting VTA^DA^ neurons (Figure S3A-B) were less excitable than BLA-projecting neurons of saline-treated mice (Figure S3C-D), as reported previously(5,8). Excitability was significantly increased in either lAcbSh-projecting or BLA-projecting VTA^DA^ neurons of morphine treated mice where slices underwent cellular withdrawal. However, in VTA slices kept in morphine, the increased excitability of either projection was not apparent (Figure S3A-D). We observed an increase in I*_h_*current in BLA-projecting (Figure S3F) but not lAcbSh-projecting VTA^DA^ neurons after morphine treatment (Figure S3E), regardless of cellular withdrawal. Further, in BLA-projecting VTA^DA^ neurons there was no difference in GIRK currents between saline- and morphine-treated mice (Figure S3G-H). These data suggest that there are projection specific differences in the response of VTA^DA^ neurons to morphine treatment. Because the morphine-induced shift in effect of endogenous LH_ox/dyn_ on VTA^DA^ neuronal firing occurred only in the BLA projection, we targeted this projection to better understand the mechanism.

### LH dynorphin release increases with repeated morphine exposure

The number of orexin-positive neurons increases in mice receiving 2 weeks of morphine injections, and in postmortem samples from former human heroin users compared to age-matched non-users(21). Based on this and other findings(20), an orexin reserve is hypothesized to increase with chronic substance use and may underlie the increased drive for opioid seeking(22). However, the number of orexin-positive neurons, fluorescence area, and integrated density within defined LH regions did not differ between 5 days of saline-and morphine-treated mice (Figures S4A–E), consistent with previous work showing that 14, but not 7 days of escalating doses of morphine increases orexin cell number(45). Next, we assessed whether peptide release was altered (Figure 3A) (37). Optical stimulation (30 Hz, 30 s) of LH_ox/dyn_ terminals in the VTA significantly increased orexin-A concentration in ChR2-expressing LH_ox/dyn_ fibres compared to mCherry-expressing controls (Figure 3A), confirming activity-dependent peptide release. However, orexin-A release in the VTA did not differ between saline- and morphine-treated mice (Figure 3B), indicating that repeated morphine exposure does not modify orexin release from LH_ox/dyn_ inputs to the VTA.

**Figure 3.**
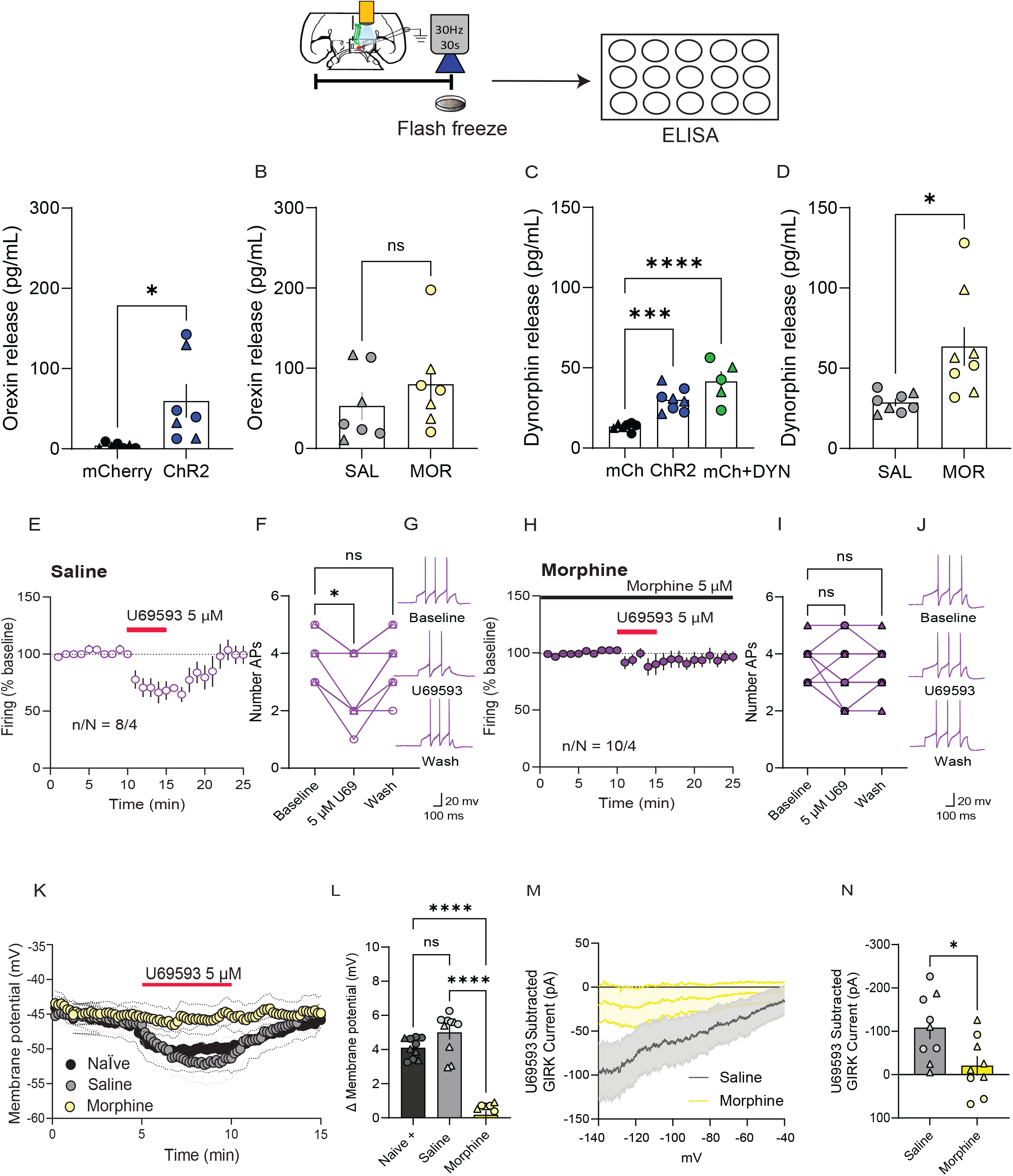
Repeated morphine exposure increases LH dynorphin release in the VTA. (A) Orexin-A concentration in VTA slice supernatants is significantly increased in ChR2-(N=7) compared to mCherry-expressing mice (N=7) following 30 Hz, 30 s optical stimulation of LH_ox/dyn_ inputs (Students t-test: t_(12)_ = 2.724, p=0.0006). (B) Orexin-A concentration in VTA slice supernatants from ChR2-expressing saline (N=7) and morphine-treated mice (N=7) is not different following 30 Hz, 30 s optical stimulation of LH_ox/dyn_ inputs (Students t-test: t_(12)_ = 0.97, p=0.35). (C) Dynorphin-A (1–17) concentration in VTA slice supernatants is increased in ChR2-expressing (n=8) compared to mCherry-expressing mice (N=7) following 30 Hz, 30 s optical stimulation of LH_ox/dyn_ inputs (P=0.0008). Addition of dynorphin-A (1-17, 140 pmol) to the supernatant of VTA slices (N=5) as a positive control is also increased compared to mCherry-expressing mice receiving optical stimulation of LH_ox/dyn_ inputs. (One way ANOVA: F_(2, 17)_ =19.55, P<0.0001) (D) Dynorphin-A (1–17) concentration in VTA slice supernatants from ChR2-expressing is greater in morphine-treated (N=8) compared to saline-treated mice (N=8) following 30 Hz, 30 s optical stimulation of LH_ox/dyn_ inputs (Student’s t-test: t_(14)_ = 2.93, p=0.01). A Rout test (Q=1%) detected one outlier in the morphine group which was removed. (E) Time course of evoked firing before and after bath application of the KOR agonist, U69593 in BLA-projecting VTA^DA^ neurons from saline-treated mice (n/N=8/4). (F) In saline-treated mice, U69593 reversibly inhibited the number of action potentials of individual BLA-projecting VTA^DA^ neurons (RM one way ANOVA: F_(2, 14)_ = 14.54, p=0.025). (G) Sample traces of evoked action potentials. (H) Time course of evoked firing before and after bath application of U69593 in BLA-projecting VTA^DA^ neurons from morphine-treated mice, in the presence of morphine in the recording solution (n/N=10/4). (I) In morphine-treated mice, bath application of U69593 did not change the number of action potentials in individual BLA-projecting VTA^DA^ neurons maintained in morphine (n/N=10/4, RM one way ANOVA: F_(2, 18)_= 1.39, p=0.53). (J) Sample traces of evoked action potentials. (K) Time course of membrane potential in response to U69593 applied to BLA-projecting VTA^DA^ neurons from saline-treated (n/N=9/4), morphine-treated (n/N=8/4), and naïve mice (n/N=9/4). (L) Maximum change in membrane potential by U69593 at 500s was significantly less in BLA-projecting VTA^DA^ neurons from morphine-treated mice compared to saline-treated or naïve mice (one way ANOVA: F_(2, 27)_ = 90.48, P<0.0001). (M) Subtracted GIRK current induced by U69593 from BLA-projecting VTA^DA^ neurons from saline- and morphine-treated mice (n/N=9/5). (N) Maximum GIRK current induced by U69593 is greater in BLA-projecting VTA^DA^ neurons from saline-treated compared to morphine-treated mice (Student’s t-test: t_(16)_= 2.63, P=0.018). Data presented as mean ± SEM or individual values from male (circles) or females (triangles). ∗p < 0.05, ∗∗∗p < 0.001.

Opioid administration is known to increase dynorphin(46–49). Optical stimulation of LH_ox/dyn_ terminals significantly increased VTA dynorphin-A 1-17 release in ChR2-expressing slices relative to mCherry controls (Figure 3C). Moreover, optically evoked dynorphin levels were similar to that of exogenous dynorphin-A 1-17 (140 nM) added to the supernatant, confirming the sensitivity of the assay. Optical stimulation of dynorphin release was significantly higher in VTA supernatants from morphine-treated mice compared to saline controls (Figure 3D). These findings indicate that repeated morphine enhances dynorphin release from LH_ox/dyn_ projections to the VTA, despite unchanged orexin release. This result seemed counterintuitive, given the inhibitory effects of dynorphin on VTA^DA^ neuronal firing and the shift to increased excitability of the BLA-projecting VTA^DA^ neurons in response to LH_ox/dyn_ optical stimulation. Therefore, we next hypothesized that elevated dynorphin levels may be desensitizing KOR receptors on BLA-projecting VTA^DA^ neurons.

### KORs on BLA-projecting VTA^DA^ neurons are uncoupled following escalating morphine treatment

We identified a saturating dose of the KOR agonist U69593 in naïve mice (5 μM; Figure S5), and found that it robustly and reversibly decreased firing of BLA-projecting VTA^DA^ neurons in saline-treated mice (Figure 3E–G). This effect was lost in morphine-treated mice (Figures S5H–J). Consistent with this, U69593 induced a strong hyperpolarization of membrane potential in BLA-projecting VTA^DA^ neurons of naïve and saline-, but not morphine-treated mice (Figure 5K, L). Current–voltage (I–V) analyses further demonstrated a reduced U69593-evoked G-protein coupled inwardly rectifying K+ (GIRK) current in morphine-treated compared with saline-treated neurons (Figure 5M, N). We next compared expression of *Oprk1* or *Hcrt1* on BLA-projecting VTA^DA^ neurons and lAcbSh-projecting VTA^DA^ neurons, using RNAscope (Figure S6). Expression of Oprk1 was similar in BLA-and lAcbSh projecting VTA^DA^ neurons. However, the number puncta expressing *Hcrt1* mRNA was significantly less than *Oprk1* mRNA in both projections (Figure S6). Together, these results show that repeated morphine exposure impairs KOR signaling on BLA-projecting VTA^DA^ neurons, eliminating the inhibitory effect of U69593 on KOR activation. KOR desensitization may reshape the orexin– dynorphin balance within the VTA and contribute to the altered modulation of BLA-projecting VTA^DA^ neurons observed after morphine treatment.

### Aticaprant prevents KOR desensitization on BLA-projecting VTA^DA^ neurons after repeated morphine treatment

Because repeated morphine desensitized KOR signaling on BLA-projecting VTA^DA^ neurons, we next asked whether co-administration of the selective, reversible KOR antagonist, aticaprant, could prevent this effect. Mice expressing ChR2 in LH_ox/dyn_ neurons and retrobeads in the BLA received the same morphine regimen with twice- daily aticaprant (Figure 4A-B). In saline-treated mice given aticaprant, optical stimulation of LH_ox/dyn_ inputs in the VTA primarily decreased firing of BLA-projecting VTA^DA^ neurons (Figure 4C–F), similar to that demonstrated in saline-treated mice in Figure 2. In mice given morphine and aticaprant, LH_ox/dyn_ optical stimulation also produced predominantly inhibitory effects on BLA-projecting VTA^DA^ neurons (Figures 4G–J), in contrast to the excitatory shift seen after morphine alone (Figure 2). Thus, aticaprant preserved KOR-mediated inhibition. Consistent with this, bath application of U69593 (5 μM) significantly reduced firing in BLA-projecting VTA^DA^ neurons from both saline + aticaprant (Figure 4K,L) and morphine + aticaprant groups (Figures 4M,N). Together, these findings demonstrate that aticaprant, a selective and reversible KOR antagonist, prevents morphine-induced KOR desensitization on BLA-projecting VTA^DA^ neurons.

**Figure 4.**
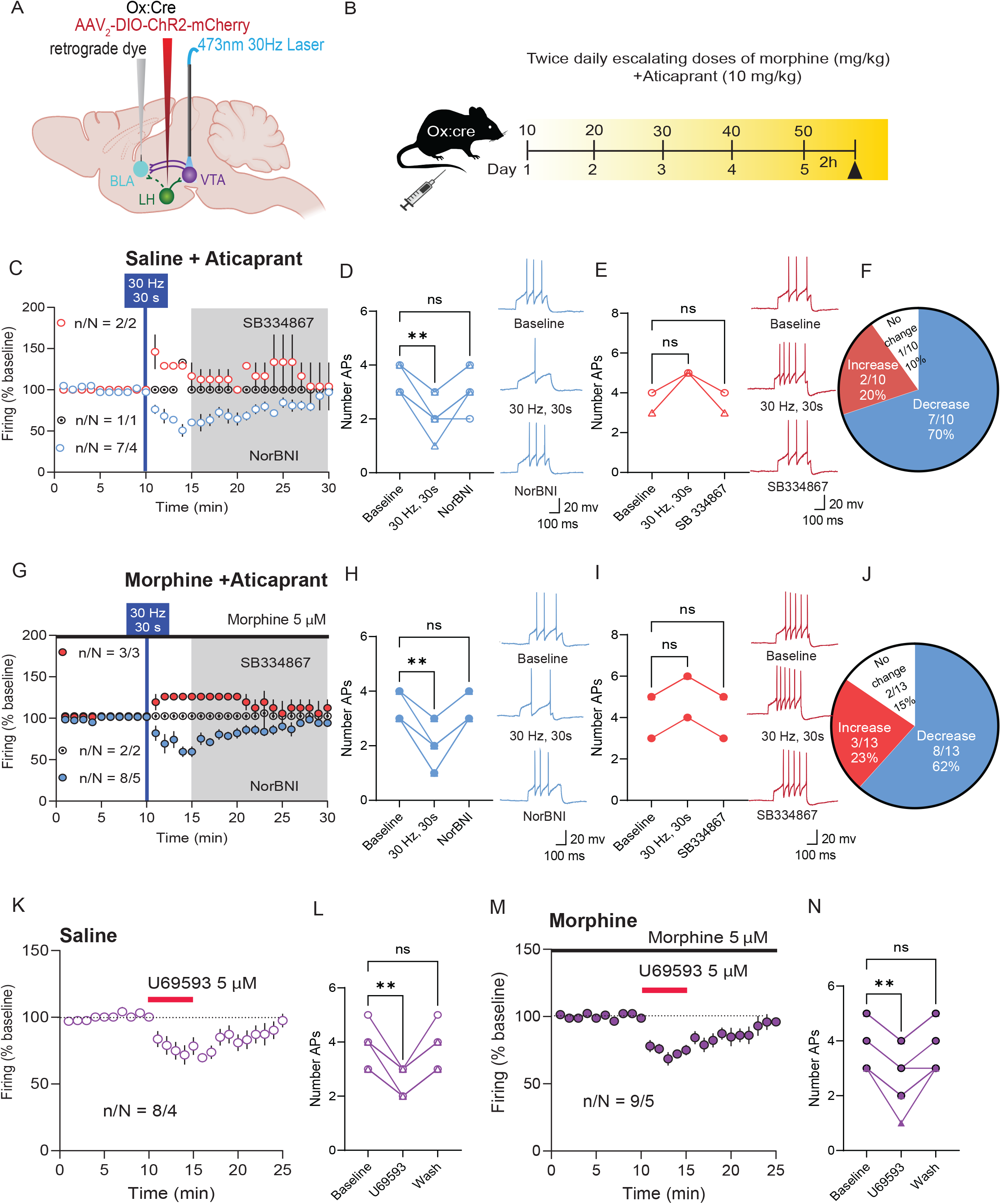
Aticaprant prevents KOR desensitization on BLA-projecting VTA^DA^ neurons after repeated morphine. (A) Schematic of viral strategy. (B) Timeline of escalating morphine treatment with twice daily aticaprant (except for day 5), a reversible KOR antagonist. (C) Time course of evoked firing before and after 30 Hz, 30s photoactivation of LH_ox/dyn_ inputs in BLA-projecting VTA^DA^ neurons from saline-treated mice given aticaprant (n/N=10/7). (D,E) Number of action potentials and sample traces during baseline, after 30 Hz, 30 s optical stimulation, and after NorBNI or SB334867 application in BLA-projecting VTA^DA^ neurons that showed more decreased responses (D; n/N=7/4, RM one-way ANOVA: F_(2,12)_ = 23.65, p=0.0066) than increased (E; n/N=2/2, F_(2, 2)_ = 9, p=0.2672) responses in BLA-projecting VTA^DA^ neurons of saline and aticaprant-treated mice. (F) Optical stimulation of LH_ox/dyn_ inputs in the VTA mostly decreases firing rate of BLA-projecting VTA^DA^ neurons in mice treated with saline + aticaprant. (G) Time course of evoked firing before and after 30 Hz, 30s photoactivation of LH_ox/dyn_ inputs in BLA-projecting VTA^DA^ neurons of morphine and aticaprant-treated mice (n/N=10/7). (H,I) Number of action potentials and sample traces during baseline, after 30 Hz, 30 s optical stimulation, and after NorBNI or SB334867 application in BLA-projecting VTA^DA^ neurons showed mostly decreased (H; n/N=8/5, RM one-way ANOVA: F_(2, 14)_=45.71, p=0.0036) or but few increased (I; n/N=3/3, p=0.13) responses in morphine and aticaprant treated mice. (J) In mice co-treated with morphine and aticaprant, the proportion of responses of BLA-projecting VTA^DA^ neurons to optical stimulation of LH_ox/dyn_ inputs in the VTA is not different from saline-and aticaprant-treated mice (χ²=0.067, p=0.79). (K) Time course of evoked firing before and after bath application of U69593 in BLA-projecting VTA^DA^ neurons from mice treated with saline + aticaprant (n/N=8/4). (L) U69593 suppresses the number of action potentials of BLA-projecting VTA^DA^ neurons (n/N=8/4, RM one-way ANOVA: F_(2, 14)_ =30.33, p=0.0036). (M) Time course of evoked firing before and after bath application of U69593 in BLA-projecting VTA^DA^ neurons from mice treated with morphine + aticaprant (n/N=8/4). (N) U69593 suppresses the number of action potentials of BLA-projecting VTA^DA^ neurons from mice treated with morphine + aticaprant (n/N=9/5, RM one-way ANOVA: F_(2,12)_ = 42.82, p=0.0029). Data presented as mean ± SEM or individual values from male (circles) or females (triangles). ∗p < 0.05, ∗∗p < 0.01.

Having established a projection-specific reduction in KOR function in BLA-projecting VTA^DA^ neurons *ex vivo*, we next asked whether this cellular adaptation translated into altered downstream dopamine output *in vivo*. We therefore combined optogenetic activation of LH_ox/dyn_ terminals the VTA with GRAB_DA_ fiber photometry in the BLA to determine the functional consequence of this adaptation within the intact LH– VTA–BLA circuit (Figure 5A). Optical stimulation of LH_ox/dyn_ inputs to the VTA produced a robust dopamine response in the BLA at each 30 Hz, 30 s stimulation (Figure 5B, C). In contrast to SB334867, administration of aticaprant profoundly increased LH_ox/dyn_ evoked BLA^DA^ (Figure 5D, E). Thus, not only does LH_ox/dyn_ activation release BLA^DA^, but this is also strongly regulated by dynorphin signaling in the VTA in naïve mice. Therefore, we predicted that because KORs are desensitized in the VTA-BLA projection in morphine-dependent mice, BLA^DA^ would be significantly increased. Indeed, BLA^DA^ evoked by optical stimulation of LH_ox/dyn_ inputs to the VTA was significantly greater in morphine-dependent mice compared to saline-treated mice (Figure 5F,G). Taken together, opioid-dependence leads to a diminished KOR regulation of BLA-projecting VTA^DA^ neurons resulting in enhanced BLA^DA^.

**Figure 5.**
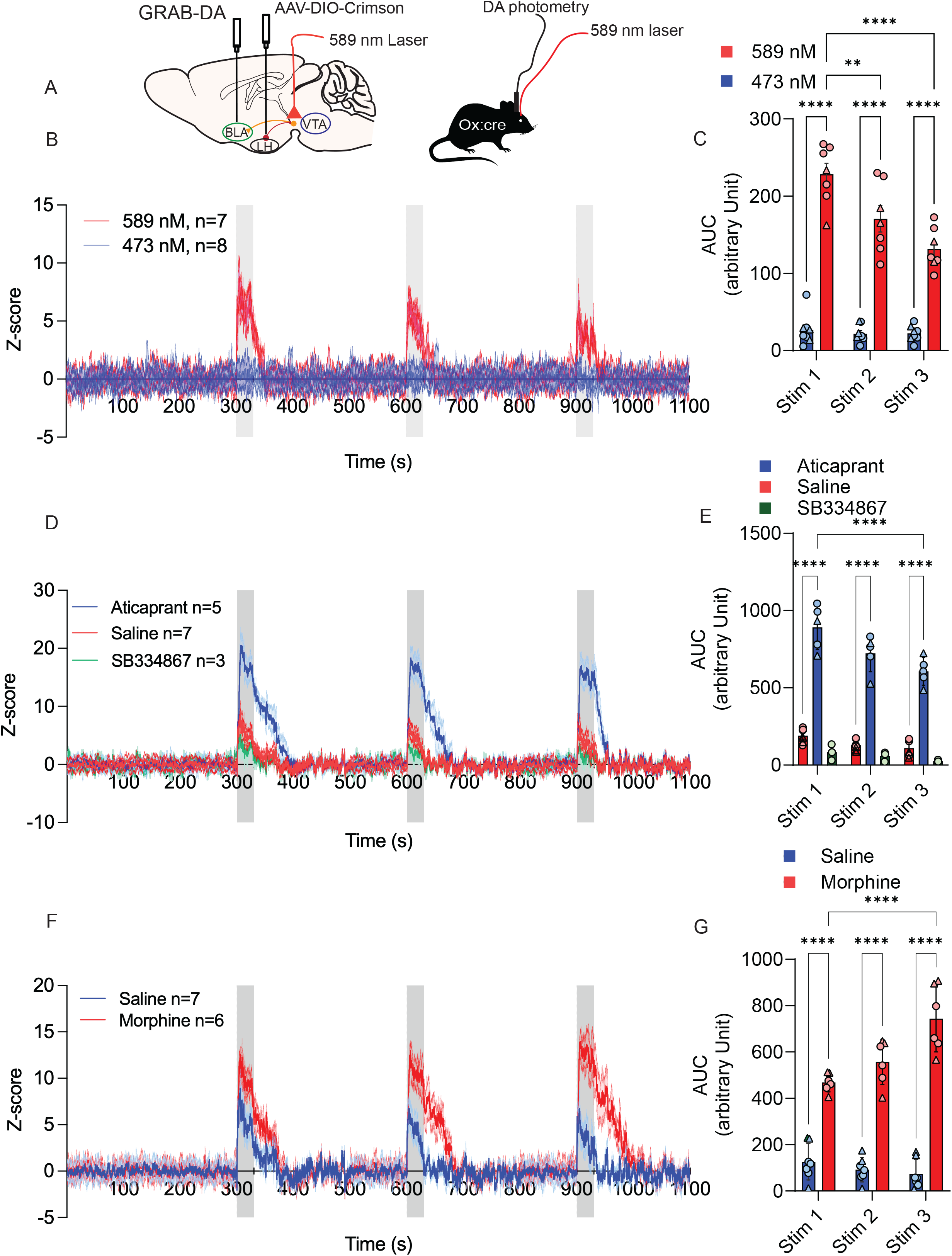
Morphine-treatment increases BLA dopamine in response to optical stimulation of LH_ox/dyn_ inputs to the VTA. (A) Schematic of viral strategy whereby GRAB_DA_ and a photometry ferrule is expressed in the BLA, chrimson is expressed in LH_ox/dyn_ neurons, and the optic ferrule is implanted to the VTA. (B) Time course of Z-scored photometry responses measuring BLA dopamine to 3 optical stimulations at 30 Hz, 30 s laser stimulation of the active (589 nM; N=7) and inactive (473 nM; N=8) laser. (C) BLA dopamine is strongly evoked by activating chrimson expressing LH_ox/dyn_ terminals in the VTA. RM two-way ANOVA (stimulation × laser wavelength interaction: F_(2,39)_ = 10.37, P = 0.0002; main effect of laser wavelength: F_(1,39)_ = 344.0, P < 0.0001; main effect of stimulation: F_(2,39)_ = 12.71, P < 0.0001). Sidak’s posthoc test showed that 589-nm stimulation evoked significantly greater AUC responses than 473-nm stimulation across all three stimulation events (P < 0.0001). There was also a decay in evoked BLA^DA^ with repeated LH_ox/dyn_ stimulation (P<0.0001). (D) Time course of Z-scored photometry responses measuring BLA dopamine to 3 optical stimulations at 30 Hz, 30 s laser stimulation in the presence of saline (red), aticaprant (blue), and SB334867 (green). (E) BLA dopamine is evoked by activating crimson-expressing LH_ox/dyn_ terminals in the VTA of saline-treated mice. This response is significantly potentiated by the KOR antagonist aticaprant, while the Ox1R antagonist SB334867 does not significantly alter dopamine release. RM two-way ANOVA (stimulation × drug interaction: F_(4,36)_ = 3.21, P = 0.024; main effect of drug: F_(2,36)_= 11.38, P = 0.0001; main effect of stimulation: F_(2,36)_ = 333.3, P < 0.0001). Sidak’s posthoc test revealed that aticaprant produced significantly greater AUC responses than saline or SB334867 across stimulation events (P < 0.0001), whereas SB334867 did not differ from saline (Stim 1: P = 0.83; Stim 2: P = 0.99; Stim 3: P = 0.99). (F) Time course of Z-scored photometry responses to 3 optical stimulations at 30 Hz, 30 s laser stimulation in morphine-treated (red; N = 6) versus saline-treated (blue; N = 7) mice. (G) Photoactivating crimson-expressing LH_ox/dyn_ terminals in the VTA of morphine-treated mice (N =6) evokes greater BLA dopamine that in saline-treated mice (N=7). RM two-way ANOVA (drug × stimulation interaction: F_(2,33)_ = 12.41, P < 0.0001; main effect of drug: F_(1,33)_ = 328.6, P < 0.0001; main effect of stimulation: F_(2,33)_ = 6.07, P = 0.0057). Sidak’s posthoc test revealed that morphine-treated mice showed significantly higher AUC values than saline controls across all stimulation events (P < 0.0001). Data presented as mean ± SEM or individual values from male (circles) or females (triangles). ∗∗p < 0.01, ∗∗∗p < 0.001, ∗∗∗p < 0.0001.

To test if BLA^DA^ was regulated by a simultaneous morphine injection, a cohort of mice treated with twice-daily morphine for 5 days were given an additional 50 mg/kg injection of morphine on the 6^th^ day during photometry recordings (Figure S7A). Morphine robustly increased BLA^DA^ and this effect was reversed by naloxone administration (Figure S7B-C). Thus, morphine significantly increases BLA^DA^ in morphine dependent mice.

### LH_ox/dyn_ modulation of BLA-projecting VTA^DA^ neurons is excitatory in fentanyl self-administering mice

Fentanyl has superseded other opioids in the opioid use landscape(50). Therefore, we developed an oral fentanyl self-administration model to test the effects of LH_ox/dyn_ stimulation on BLA-projecting VTA^DA^ neurons (Figure 6A, B). Mice acquired fixed ratio 1 (FR1) fentanyl self-administration and discriminated between active and inactive levers showing stable fentanyl consumption across 20 sessions (Figure S8A–E). Sucrose self-administering mice showed similar task acquisition on an FR1 schedule (Figure S8F–J). Oral fentanyl self-administration produced opioid dependence as mice administered naloxone following the final self-administration session displayed significantly greater withdrawal signs compared to sucrose self-administering mice receiving naloxone (Figure S9). In sucrose self-administering mice, optical stimulation of LH_ox/dyn_ inputs in the VTA primarily decreased firing of BLA-projecting VTA^DA^ neurons (Figure 6C–F).

**Figure 6.**
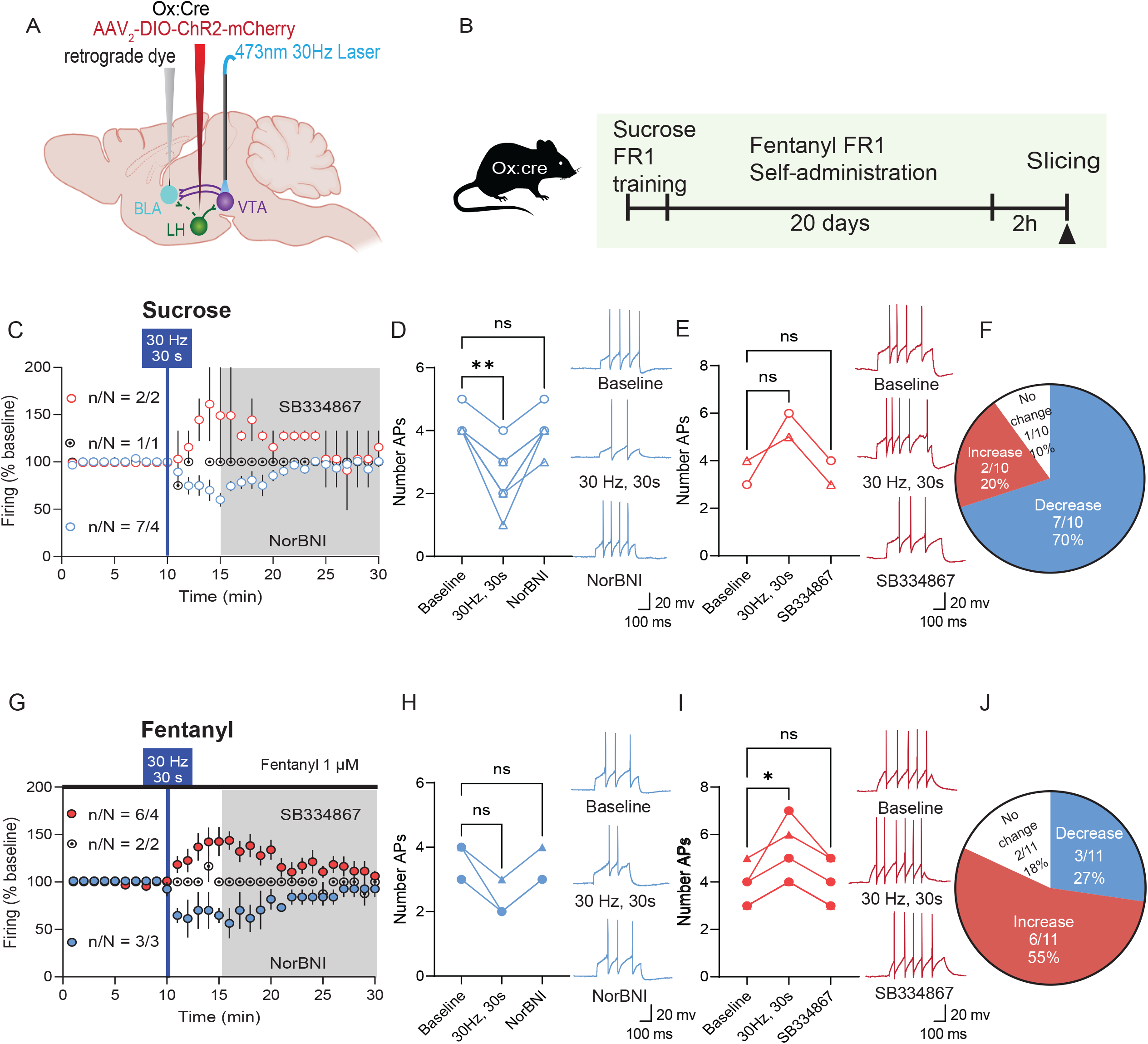
Voluntary oral-fentanyl self-administration increased excitability of BLA-projecting VTA^DA^ neurons in response to optical stimulation of LH_ox/dyn_ inputs compared to sucrose self-administering mice. (A) Schematic of viral strategy. (B) Experimental timeline of FR1 fentanyl self-administration training and subsequent electrophysiological recordings. (C) Time course of evoked firing before and after 30 Hz, 30s photoactivation of LH_ox/dyn_ inputs in BLA-projecting VTA^DA^ neurons of sucrose mice (n/N=10/7). (D-E) Number of action potentials and sample traces during baseline, after 30 Hz, 30 s optical stimulation, and after NorBNI or SB334867 application in BLA-projecting VTA^DA^ neurons were decreased (D; n/N=7/4, RM one-way ANOVA: F_(2, 12)_=28.50, p=0.0066) but rarely increased (E; n/N=2/2, RM one-way ANOVA: F_(2, 2)_=4, p=0.27) to LH_ox/dyn_ optical stimulation in sucrose self-administering mice. (F) Optical stimulation of LH_ox/dyn_ inputs in the VTA mostly decreases firing responses of BLA-projecting VTA^DA^ neurons in sucrose self-administering mice. (G) Time course of evoked firing before and after 30 Hz, 30s photoactivation of LH_ox/dyn_ inputs in BLA-projecting VTA^DA^ neurons of fentanyl self-administering mice (n/N=11/9), recorded in the presence of fentanyl (1 μM). (H-I) Number of action potentials and sample traces during baseline, after 30 Hz, 30 s optical stimulation, and after NorBNI or SB334867 application in BLA-projecting VTA^DA^ neurons were rarely decreased (H; n/N=3/3, RM one-way ANOVA: F_(2, 14)_=13, p=0.082) but mostly increased (I; n/N=6/4, RM one-way ANOVA: F_(2, 10)_= 19, p=0.012) in response to LH_ox/dyn_ optical stimulation in fentanyl self-administering mice. (J) Optical stimulation of LH_ox/dyn_ inputs in the VTA mostly increases firing rate of BLA-projecting VTA^DA^ neurons in fentanyl self-administering mice compared to sucrose self-administering mice (χ²=4.34, p=0.037). Data presented as mean ± SEM or individual values from male (circles) or females (triangles). ∗p < 0.05, ∗∗p < 0.01.

Inhibitory responses were washed off by the KOR antagonist, NorBNI (Figure 6D). In contrast, in mice that self-administered fentanyl, optical stimulation of LH_ox/dyn_ terminals predominantly increased firing of BLA-projecting VTA^DA^ neurons (Figure 6G–J). This excitatory effect is similar to the effect observed after experimenter-administered morphine (Figure 2), suggesting that chronic opioid exposure, regardless of administration route, produces a similar shift in LH_ox/dyn_ modulation. Together, these findings indicate that voluntary fentanyl intake also shifts the inhibitory LH_ox/dyn_ influence on BLA-projecting VTA^DA^ neurons into an excitatory neuromodulatory effect.

## Discussion

We found that opioids shift the neuromodulatory capacity of the LH_ox/dyn_ input to BLA-projecting VTA^DA^ neurons from inhibitory to excitatory. This is driven by increased dynorphin release resulting in KOR desensitization on BLA-projecting VTA^DA^ neurons. These results reveal an opioid-induced shift in peptide co-transmission within the LH– VTA–BLA circuit, demonstrating distinct modulatory roles for orexin and dynorphin in the VTA output and suggesting a mechanism by which chronic opioid exposure can increase BLA^DA^. Given that BLA^DA^ responds to predictive cues and reward-identity memories (51,52), we provide a potential mechanism for how repeated opioid use increases reinforcement of opioid-related cues and behaviours.

Under physiological conditions, orexin and dynorphin act as a counterbalancing state signal that gates dopamine information flow to regions including the BLA and the lAcbSh. Given that orexin and dynorphin are co-released into the VTA, they generate a state change signal that may differentially gate information to different VTA projections. Optical stimulation of the LH_ox/dyn_ input to the VTA predominantly inhibits BLA-projecting VTA^DA^ neurons, due to dynorphin/KOR-mediated effects that dominate over orexin/Ox1R, while producing bidirectional responses in lAcbSh- and medial AcbSh-projecting neurons(14). Thus, state-dependent peptide release may balance ensembles of VTA^DA^ neurons within different projections to influence their final output. Repeated opioids disrupt this process and redirects VTA^DA^ activity to drive different circuits. While modulation of the projection to the lAcbSh is unchanged, activity of the BLA-projecting VTA^DA^ neurons is enhanced leading to increased BLA^DA^ release. Consistent with our findings, intersectional c-Fos labeling in BLA-projecting VTA^DA^ neurons is more predominant than lAcbSh-projecting VTA^DA^ neurons after repeated opioids(32).

KORs are expressed both presynaptically(56) and postsynaptically on dopamine neuron somata in both BLA- and NAc-projections(58), where their activation inhibits neuronal firing (25,26,57). Our recordings were in the presence of synaptic blockers, excluding effects of presynaptic KORs (56). Notably, we found that KORs are highly expressed on BLA-projecting VTA^DA^ neurons, and to a greater extent than Ox1R expression in this projection, likely underlying why this VTA^DA^ projection is highly sensitive to inhibitory regulation of LH dynorphin(14).

Enhanced excitatory neuromodulation of the LH_ox/dyn_ input to BLA-projecting VTA^DA^ resulted from decreased KOR function. After repeated morphine, a maximal concentration of a KOR agonist no longer suppressed firing or activated GIRK currents of BLA-projecting VTA^DA^ neurons. These data suggest that whilst there is an increased dynorphin release in the VTA, KOR function is decreased in BLA-projecting VTA^DA^ neurons resulting in increased BLA^DA^, presumably driving reward-cue encoding(52). While these findings contradict a prevailing hypothesis that there in an increase in KOR tone leading to an increased ‘brake’ which opposes reward processing(54), our results are supported by findings that stress-induced dynorphin release mediates the increased rewarding effect of cocaine(59–61). Prior activation of KOR via an agonist or stress-induced dynorphin release caused a potentiation of cocaine conditioned place preference, that was blocked by inhibition of KORs or gene deletion of dynorphin(61). Prolonged or continuous exposure to elevated dynorphin may result in receptor desensitization or a change in functional effector coupling, shifting the balance toward orexin-driven excitation, at least in the BLA projection. Consistent with our results showing that chronic morphine increases dynorphin release in the VTA from optically stimulated LH_ox/dyn_ inputs, chronic morphine increases prodynorphin gene expression in mice (55) and dynorphin-A and B peptides in the VTA and NAc of Fisher rats(47). Furthermore, sustained dynorphin exposure can reduce receptor affinity, without altering KORs density, leading to KOR desensitization(37,62)

Several hypotheses may explain why after repeated morphine, KORs no longer hyperpolarize VTA^DA^ neurons in the BLA projection. First, chronic morphine decreases gene expression of certain K+ channels in the VTA, such as *kcnab2*, *kcnj2* and *girk3*, and these adaptations are associated with increased excitability of VTA^DA^ neurons(63). Mice repeatedly exposed to methamphetamine had decreased GABA_B_ and D2 receptor activated GIRK currents(64). Thus, impaired KOR-mediated inhibition of firing and GIRK currents observed in morphine-treated mice could be due to heterologous mechanisms where GIRK is downregulated. However, this mechanism is unlikely as we did not observe differences in basal GIRK currents in BLA-projecting VTA^DA^ neurons of saline or morphine treated mice, suggesting that morphine treatment induces an uncoupling of KOR and its effectors rather than decreased GIRK expression in BLA-projecting VTA^DA^ neurons. Secondly, although dynorphins act on KORs, their metabolic products are potent ligands at other opioid receptors. Dynorphin fragments can be cleaved into leu-enkephalin, which have nanomolar affinity for MOR or δ-opioid receptors(65–67). Such interactions raise the possibility that dynorphin’s net effect in the LH _ox/dyn_ → VTA^DA^ pathway may be mediated partly through MOR or δ-opioid receptors rather than solely through KORs. The selective KOR agonist did not decrease firing or alter GIRK currents in the VTA of morphine-exposed mice, suggesting that these effects were selective to modulation of KORs rather than other opioid receptors. Furthermore, MORs are primarily presynaptically located within the VTA(43,68) and all recordings were in the presence of synaptic blockers. However, there is a small proportion of VGlut2 expressing VTA neurons that also express MOR and decrease glutamate release in response to opioid agonists (69,70). Local MOR+ VGlut2 neurons synapse onto VTA^DA^ neurons and can drive dopamine release in the NAc core, an effect that is reduced by oxycodone (70). It is unknown whether MOR+ VGlut2 neurons project directly to the BLA or modulate BLA-projecting VTA^DA^ neurons. Given that acute morphine administration increased BLA^DA^, even after 5 days of escalating morphine doses, it is unlikely that this effect was mediated by MOR on VGlut2 neurons. Thirdly, an increase in excitability in morphine-treated mice may be due to KOR-induced increase hyperpolarization activated cation current (*I*_h_). In mice exposed to footshock stress or application of corticotropin releasing factor to VTA slices, KOR agonists depolarized a subpopulation of VTA^DA^ neurons and attenuated conditioned place aversion via increased KOR coupling to *I*_h_ (71). This is consistent with our observation that morphine treatment increases I*_h_* expression in BLA-projecting VTA^DA^ neurons. Notably, while KORs are classically coupled to inhibitory G_i_ proteins, evidence indicates that under certain conditions they may couple to stimulatory G_s_ proteins, as observed in dorsal root ganglia(72,73).

Why KORs on BLA but not lAcbSh-projecting neurons are recruited for desensitization by morphine is unclear. KOR activation can recruit MEK/ERK activation (74,75) and MAPK signaling cascades, in particular p38 (76–78). Sustained KOR activation of the p38 signaling cascade can produce KOR internalization via β-arrestin signaling (77,79). Thus, it is possible that p38 levels may be differentially expressed in VTA^DA^ neurons projecting to the BLA compared to those projecting to the lAcbSh. Consistent with this, even though p38 is expressed in the VTA, p38a MAPK was not required for KOR inhibition of evoked NAc dopamine (80).

The selective recruitment of the VTA→BLA pathway has important implications for addiction-related behavior. Firstly, we report that BLA^DA^ is modulated by LH_ox/dyn_ release in the VTA. This effect is greater in the presence of KOR antagonists, suggesting that under physiological conditions, KOR activation in the VTA serves as a brake on BLA^DA^. BLA^DA^ induced by photostimulation of LH_ox/dyn_ inputs in the VTA was not modulated by Ox1R signaling, consistent with *ex vivo* firing recordings. After repeated morphine, BLA^DA^ induced by photostimulation of LH_ox/dyn_ inputs in the VTA was increased, consistent with reduced KOR signaling in this pathway. GRAB_DA_ photometry allowed us to determine whether the projection-specific loss of KOR-mediated inhibition identified electrophysiologically was expressed as altered BLA^DA^ output in the intact circuit. Importantly, pharmacological manipulation during these experiments provided additional mechanistic support: KOR antagonism markedly potentiated LH_ox/dyn_-evoked BLA^DA^, whereas Ox1R antagonism produced no detectable change. Together, the in vivo and ex vivo approaches provide convergent evidence that reduced KOR-mediated inhibitory control enhances LH_ox/dyn_-driven dopamine output to the BLA following repeated opioid exposure. Moreover, morphine robustly increases BLA^DA^ in opioid-dependent mice, suggesting that MOR signaling in this pathway is functional. Dopaminergic signaling in the BLA plays a critical role in salience attribution, emotional learning, fear conditioning, and the formation of cue–outcome associations(51,81–83). Because repeated morphine exposure or fentanyl self-administration shifts LH_ox/dyn_ modulation of this pathway toward excitation, the emotional and memory encoding of drug-associated cues and contexts may be enhanced, consistent with models of incentive sensitization and maladaptive reward learning(84,85). Future studies should test if this could contribute to the persistence of craving, heightened responsivity to drug-paired stimuli, and vulnerability to relapse long after drug cessation(86–88).

In summary, we find that chronic opioid treatment shifts state-dependent LH peptide release to the VTA which redistributes ensemble activity of VTA^DA^ neurons to enhance dopamine output in the BLA. We demonstrate that this occurs in both experimenter-administered morphine and voluntary oral fentanyl self-administration models of opioid dependence. Further, we demonstrate that opioids robustly increase BLA^DA^ and that LH_ox/dyn_ inputs to the VTA regulate BLA^DA^ more strongly in morphine-dependent mice, due to an uncoupling of KOR signaling. Together our results suggest that while dynorphin peptide release is upregulated, KOR activity no longer hyperpolarizes BLA-projecting VTA^DA^ neurons after repeated morphine. This work provides a new perspective to the prevailing ‘KOR-brake’ model. Some projections may be differently regulated by KORs after chronic opioids, as this state-dependent shift applied selectively to the BLA projection, while LH_ox/dyn_ regulation of the lAcbSh projection remained intact. Future studies should investigate whether opioids alter LH_ox/dyn_ neuromodulation of other dopamine projections such as the medial AcbSh or mPFC, which have distinct roles in reward-related processing.

## Acknowledgements

The authors would like to acknowledge the Hotchkiss Brain Institute advanced microscopy facility. This research was performed at the University of Calgary which is located on the unceded traditional territories of the people of the Treaty 7 region in Southern Alberta, which includes the Blackfoot Confederacy (including the Siksika, Piikuni, Kainai First Nations), the Tsuut’ina, and the Stoney Nakoda (including the Chiniki, Bearspaw, and Goodstoney First Nations). The City of Calgary is also home to Metis Nation of Alberta.

## Funding

This work was supported by an CIHR project grant (PJT-540588 to S.L.B.) and a Canada Research Chair (950-232211). A.M. was supported by a Mathison Centre for Research and Education Postdoctoral Fellowship.

## Author Contributions

AM performed surgeries and electrophysiological experiments. HS performed surgeries and photometry experiments. II performed withdrawal precipitation and scoring. AM, HS, II performed oral fentanyl self-administration experiments. MQ performed immunohistochemistry and imaging experiments. AM, HS, II, MQ, and SLB analyzed the data. AM and SLB wrote first drafts of the manuscript. AM, HS, II, MQ, and SLB revised the manuscript.

## Conflict of Interest Statement

The authors declare no competing financial or other conflicts of interest.

## Notes

### Competing Interest Statement

The authors have declared no competing interest.

## References

1. Opioids | Canadian Centre on Substance Use and Addiction (n.d.): Retrieved December 1, 2025, from https://www.ccsa.ca/en/guidance-tools-resources/substances/opioids?field_areas_of_expertise_target_id=10365

2. Doyle MA, Mazei-Robison MS (2021): Opioid-Induced Molecular and Cellular Plasticity of Ventral Tegmental Area Dopamine Neurons. Cold Spring Harb Perspect Med 11: a039362.

3. Nestler EJ, Lüscher C (2019): The Molecular Basis of Drug Addiction: Linking Epigenetic to Synaptic and Circuit Mechanisms. Neuron 102: 48–59.

4. Kahnt T, Schoenbaum G (2025): The curious case of dopaminergic prediction errors and learning associative information beyond value. Nat Rev Neurosci 26: 169–178.

5. Lammel S, Hetzel A, Häckel O, Jones I, Liss B, Roeper J (2008): Unique Properties of Mesoprefrontal Neurons within a Dual Mesocorticolimbic Dopamine System. Neuron 57: 760–773.

6. Beier KT, Steinberg EE, DeLoach KE, Xie S, Miyamichi K, Schwarz L, et al. (2015): Circuit Architecture of VTA Dopamine Neurons Revealed by Systematic Input-Output Mapping. Cell 162: 622–634.

7. Morel C, Montgomery SE, Li L, Durand-de Cuttoli R, Teichman EM, Juarez B, et al. (2022): Midbrain projection to the basolateral amygdala encodes anxiety-like but not depression-like behaviors. Nat Commun 13: 1532.

8. Baimel C, Lau BK, Qiao M, Borgland SL (2017): Projection-Target-Defined Effects of Orexin and Dynorphin on VTA Dopamine Neurons. Cell Reports 18: 1346–1355.

9. Peyron C, Tighe DK, van den Pol AN, de Lecea L, Heller HC, Sutcliffe JG, Kilduff TS (1998): Neurons Containing Hypocretin (Orexin) Project to Multiple Neuronal Systems. J Neurosci 18: 9996– 10015.

10. Baldo BA, Daniel RA, Berridge CW, Kelley AE (2003): Overlapping distributions of orexin/hypocretin- and dopamine-?-hydroxylase immunoreactive fibers in rat brain regions mediating arousal, motivation, and stress. J Comp Neurol 464: 220–237.

11. Fadel J, Deutch AY (2002): Anatomical substrates of orexin–dopamine interactions: lateral hypothalamic projections to the ventral tegmental area. Neuroscience 111: 379–387.

12. Korotkova TM, Sergeeva OA, Eriksson KS, Haas HL, Brown RE (2003): Excitation of Ventral Tegmental Area Dopaminergic and Nondopaminergic Neurons by Orexins/Hypocretins. J Neurosci 23: 7–11.

13. Vittoz NM, Schmeichel B, Berridge CW (2008): Hypocretin /orexin preferentially activates caudomedial ventral tegmental area dopamine neurons. European Journal of Neuroscience 28: 1629–1640.

14. Mohammadkhani A, Qiao M, Borgland SL (2024): Distinct Neuromodulatory Effects of Endogenous Orexin and Dynorphin Corelease on Projection-Defined Ventral Tegmental Dopamine Neurons. J Neurosci 44: e0682242024.

15. Borgland SL, Chang S-J, Bowers MS, Thompson JL, Vittoz N, Floresco SB, et al. (2009): Orexin A/Hypocretin-1 Selectively Promotes Motivation for Positive Reinforcers. Journal of Neuroscience 29: 11215–11225.

16. Thompson JL, Borgland SL (2011): A role for hypocretin/orexin in motivation. Behavioural Brain Research 217: 446–453.

17. Fragale JE, Pantazis CB, James MH, Aston-Jones G (2019): The role of orexin-1 receptor signaling in demand for the opioid fentanyl. Neuropsychopharmacology 44: 1690–1697.

18. Mahler SV, Aston-Jones GS (2012): Fos Activation of Selective Afferents to Ventral Tegmental Area during Cue-Induced Reinstatement of Cocaine Seeking in Rats. J Neurosci 32: 13309–13325.

19. Baimel C, Borgland SL (2015): Orexin Signaling in the VTA Gates Morphine-Induced Synaptic Plasticity. J Neurosci 35: 7295–7303.

20. James MH, Stopper CM, Zimmer BA, Koll NE, Bowrey HE, Aston-Jones G (2018): Increased Number and Activity of a Lateral Subpopulation of Hypothalamic Orexin/Hypocretin Neurons Underlies the Expression of an Addicted State in Rats. Biol Psychiatry. 10.1016/j.biopsych.2018.07.022

21. Thannickal TC, John J, Shan L, Swaab DF, Wu M-F, Ramanathan L, et al. (2018): Opiates increase the number of hypocretin-producing cells in human and mouse brain and reverse cataplexy in a mouse model of narcolepsy. Sci Transl Med 10. 10.1126/scitranslmed.aao4953

22. James MH, Aston-Jones G (2022): Orexin Reserve: A Mechanistic Framework for the Role of Orexins (Hypocretins) in Addiction. Biol Psychiatry 92: 836–844.

23. Chou TC, Lee CE, Lu J, Elmquist JK, Hara J, Willie JT, et al. (2001): Orexin (Hypocretin) Neurons Contain Dynorphin. J Neurosci 21: RC168–RC168.

24. Muschamp JW, Hollander JA, Thompson JL, Voren G, Hassinger LC, Onvani S, et al. (2014): Hypocretin (orexin) facilitates reward by attenuating the antireward effects of its cotransmitter dynorphin in ventral tegmental area. Proceedings of the National Academy of Sciences 111: E1648–E1655.

25. Margolis EB, Hjelmstad GO, Bonci A, Fields HL (2003): κ-Opioid Agonists Directly Inhibit Midbrain Dopaminergic Neurons. J Neurosci 23: 9981–9986.

26. Ford CP, Beckstead MJ, Williams JT (2007): Kappa Opioid Inhibition of Somatodendritic Dopamine Inhibitory Postsynaptic Currents. Journal of Neurophysiology 97: 883–891.

27. Cayir S, Zhornitsky S, Barzegary A, Sotomayor-Carreño E, Sarfo-Ansah W, Funaro MC, et al. (2024): A review of the kappa opioid receptor system in opioid use. Neuroscience & Biobehavioral Reviews 162: 105713.

28. Marchette RCN, Vendruscolo LF, Koob GF (2025): The Dynorphin/-Opioid Receptor System at the Interface of Hyperalgesia/Hyperkatifeia and Addiction. Curr Addict Rep 12: 11.

29. Chen M, Zhao Y, Yang H, Luan W, Song J, Cui D, et al. (2015): Morphine disinhibits glutamatergic input to VTA dopamine neurons and promotes dopamine neuron excitation. eLife 4: e09275.

30. Mohammadkhani A, Mitchell C, James MH, Borgland SL, Dayas CV (2024): Contribution of hypothalamic orexin (hypocretin) circuits to pathologies of motivation. British J Pharmacology 181: 4430–4449.

31. Georgescu D, Zachariou V, Barrot M, Mieda M, Willie JT, Eisch AJ, et al. (2003): Involvement of the Lateral Hypothalamic Peptide Orexin in Morphine Dependence and Withdrawal. J Neurosci 23: 3106–3111.

32. Morison SL, Doster LB, Vigotsky AD, Centeno MV, Lopez Gonzalez Del Rey N, Apkarian AV, Awatramani R (2025): Recruitment of specific dopamine neuron sub-circuits by opioids. Addiction Neuroscience 17: 100233.

33. Li Y, van den Pol AN (2008): Mu-opioid receptor-mediated depression of the hypothalamic hypocretin/orexin arousal system. J Neurosci 28: 2814–2819.

34. Berry EA, Huhulea EN, Ishibashi M, McGregor R, Siegel JM, Leonard CS (2025): Repeated Morphine Exposure Reversibly Impairs Spike Generation and Repetitive Firing in a Functionally Distinct Subpopulation of Orexin Neurons. J Neurosci 45: e0563252025.

35. van den Hoogen NJ, Kwok CHT, Trang T (2021): Identifying the Neurodevelopmental Differences of Opioid Withdrawal. Cell Mol Neurobiol 41: 1145–1155.

36. Burma NE, Bonin RP, Leduc-Pessah H, Baimel C, Cairncross ZF, Mousseau M, et al. (2017): Blocking microglial pannexin-1 channels alleviates morphine withdrawal in rodents. Nat Med 23: 355– 360.

37. Li Y, van den Pol AN (2006): Differential target-dependent actions of coexpressed inhibitory dynorphin and excitatory hypocretin/orexin neuropeptides. J Neurosci 26: 13037–13047.

38. Rosin DL, Weston MC, Sevigny CP, Stornetta RL, Guyenet PG (2003): Hypothalamic orexin (hypocretin) neurons express vesicular glutamate transporters VGLUT1 or VGLUT2. J Comp Neurol 465: 593–603.

39. Thomas CS, Mohammadkhani A, Rana M, Qiao M, Baimel C, Borgland SL (2022): Optogenetic stimulation of lateral hypothalamic orexin/dynorphin inputs in the ventral tegmental area potentiates mesolimbic dopamine neurotransmission and promotes reward-seeking behaviours. Neuropsychopharmacol 47: 728–740.

40. Bagley EE, Gerke MB, Vaughan CW, Hack SP, Christie MJ (2005): GABA transporter currents activated by protein kinase A excite midbrain neurons during opioid withdrawal. Neuron 45: 433–445.

41. Ingram SL, Vaughan CW, Bagley EE, Connor M, Christie MJ (1998): Enhanced Opioid Efficacy in Opioid Dependence Is Caused by an Altered Signal Transduction Pathway. J Neurosci 18: 10269–10276.

42. Mohammadkhani A, Qiao M, Borgland SL (2024): Distinct neuromodulatory effects of endogenous orexin and dynorphin corelease on projection-defined ventral tegmental dopamine neurons. J Neurosci e0682242024.

43. Matsui A, Jarvie BC, Robinson BG, Hentges ST, Williams JT (2014): Separate GABA afferents to dopamine neurons mediate acute action of opioids, development of tolerance, and expression of withdrawal. Neuron 82: 1346–1356.

44. Jalabert M, Bourdy R, Courtin J, Veinante P, Manzoni OJ, Barrot M, Georges F (2011): Neuronal circuits underlying acute morphine action on dopamine neurons. Proc Natl Acad Sci U S A 108: 16446–16450.

45. Thannickal TC, John J, Shan L, Swaab DF, Wu M-F, Ramanathan L, et al. (2018): Opiates increase the number of hypocretin-producing cells in human and mouse brain and reverse cataplexy in a mouse model of narcolepsy. Sci Transl Med 10: eaao4953.

46. Nylander I, Vlaskovska M, Terenius L (1995): The effects of morphine treatment and morphine withdrawal on the dynorphin and enkephalin systems in Sprague-Dawley rats. Psychopharmacology (Berl) 118: 391–400.

47. Nylander I, Vlaskovska M, Terenius L (1995): Brain dynorphin and enkephalin systems in Fischer and Lewis rats: effects of morphine tolerance and withdrawal. Brain Res 683: 25–35.

48. Nylander I, Stenfors C, Tan-No K, Mathé AA, Terenius L (1997): A comparison between microwave irradiation and decapitation: basal levels of dynorphin and enkephalin and the effect of chronic morphine treatment on dynorphin peptides. Neuropeptides 31: 357–365.

49. Cappendijk SL, Hurd YL, Nylander I, van Ree JM, Terenius L (1999): A heroin-, but not a cocaine-expecting, self-administration state preferentially alters endogenous brain peptides. Eur J Pharmacol 365: 175–182.

50. Skolnick P, Paavola J, Heidbreder C (2024): Synthetic opioids have disrupted conventional wisdom for treating opioid overdose. Drug and Alcohol Dependence Reports 12: 100268.

51. Lutas A, Kucukdereli H, Alturkistani O, Carty C, Sugden AU, Fernando K, et al. (2019): State-specific gating of salient cues by midbrain dopaminergic input to basal amygdala. Nat Neurosci 22: 1820–1833.

52. Sias AC, Jafar Y, Goodpaster CM, Ramírez-Armenta K, Wrenn TM, Griffin NK, et al. (2024): Dopamine projections to the basolateral amygdala drive the encoding of identity-specific reward memories. Nat Neurosci 27: 728–736.

53. Mahler SV, Moorman DE, Smith RJ, James MH, Aston-Jones G (2014): Motivational activation: a unifying hypothesis of orexin/hypocretin function. Nat Neurosci 17: 1298–1303.

54. Tejeda HA, Shippenberg TS, Henriksson R (2012): The dynorphin/κ-opioid receptor system and its role in psychiatric disorders. Cell Mol Life Sci 69: 857–896.

55. McClung CA, Nestler EJ, Zachariou V (2005): Regulation of Gene Expression by Chronic Morphine and Morphine Withdrawal in the Locus Ceruleus and Ventral Tegmental Area. J Neurosci 25: 6005– 6015.

56. Margolis EB, Hjelmstad GO, Bonci A, Fields HL (2005): Both Kappa and Mu Opioid Agonists Inhibit Glutamatergic Input to Ventral Tegmental Area Neurons. Journal of Neurophysiology 93: 3086– 3093.

57. Svingos AL, Colago EEO, Pickel VM (1999): Cellular Sites for Dynorphin Activation of κ-Opioid Receptors in the Rat Nucleus Accumbens Shell. J Neurosci 19: 1804–1813.

58. Ford CP, Mark GP, Williams JT (2006): Properties and opioid inhibition of mesolimbic dopamine neurons vary according to target location. J Neurosci 26: 2788–2797.

59. Redila VA, Chavkin C (2008): Stress-induced reinstatement of cocaine seeking is mediated by the kappa opioid system. Psychopharmacology (Berl) 200: 59–70.

60. McLaughlin JP, Land BB, Li S, Pintar JE, Chavkin C (2006): Prior Activation of Kappa Opioid Receptors by U50,488 Mimics Repeated Forced Swim Stress to Potentiate Cocaine Place Preference Conditioning [no. 4]. Neuropsychopharmacology 31: 787–794.

61. McLaughlin JP, Marton-Popovici M, Chavkin C (2003): Kappa opioid receptor antagonism and prodynorphin gene disruption block stress-induced behavioral responses. J Neurosci 23: 5674– 5683.

62. Raynor K, Kong H, Hines J, Kong G, Benovic J, Yasuda K, et al. (1994): Molecular mechanisms of agonist-induced desensitization of the cloned mouse kappa opioid receptor. The Journal of Pharmacology and Experimental Therapeutics 270: 1381–1386.

63. Mazei-Robison MS, Koo JW, Friedman AK, Lansink CS, Robison AJ, Vinish M, et al. (2011): Role for mTOR signaling and neuronal activity in morphine-induced adaptations in ventral tegmental area dopamine neurons. Neuron 72: 977–990.

64. Munoz MB, Padgett CL, Rifkin R, Terunuma M, Wickman K, Contet C, et al. (2016): A Role for the GIRK3 Subunit in Methamphetamine-Induced Attenuation of GABAB Receptor-Activated GIRK Currents in VTA Dopamine Neurons. J Neurosci 36: 3106–3114.

65. Zamir N, Weber E, Palkovits M, Brownstein M (1984): Differential processing of prodynorphin and proenkephalin in specific regions of the rat brain. Proc Natl Acad Sci USA 81: 6886–6889.

66. Zhang S, Tong Y, Tian M, Dehaven RN, Cortesburgos L, Mansson E, et al. (1998): Dynorphin A as a Potential Endogenous Ligand for Four Members of the Opioid Receptor Gene Family. The Journal of Pharmacology and Experimental Therapeutics 286: 136–141.

67. Morgan M, Heffernan A, Benhabib F, Wagner S, Hewavitharana AK, Shaw PN, Cabot PJ (2017): The efficacy of Dynorphin fragments at the κ, μ and δ opioid receptor in transfected HEK cells and in an animal model of unilateral peripheral inflammation. Peptides 89: 9–16.

68. Johnson SW, North RA (1992): Opioids excite dopamine neurons by hyperpolarization of local interneurons. J Neurosci 12: 483–488.

69. Miranda-Barrientos J, Chambers I, Mongia S, Liu B, Wang H-L, Mateo-Semidey GE, et al. (2021): Ventral tegmental area GABA, glutamate, and glutamate-GABA neurons are heterogeneous in their electrophysiological and pharmacological properties. European Journal of Neuroscience 54: 4061–4084.

70. McGovern DJ, Polter AM, Prévost ED, Ly A, McNulty CJ, Rubinstein B, Root DH (2023): Ventral tegmental area glutamate neurons establish a mu-opioid receptor gated circuit to mesolimbic dopamine neurons and regulate opioid-seeking behavior. Neuropsychopharmacol 48: 1889– 1900.

71. Margolis EB (2025): A switch in kappa opioid receptor signaling from inhibitory to excitatory induced by stress in a subset of cortically-projecting dopamine neurons. bioRxiv 2025.08.09.669424.

72. Shen K, Crain S (1994): Nerve growth factor rapidly prolongs the action potential of mature sensory ganglion neurons in culture, and this effect requires activation of Gs-coupled excitatory kappa-opioid receptors on these cells. J Neurosci 14: 5570–5579.

73. Hampson RE, Mu J, Deadwyler SA (2000): Cannabinoid and Kappa Opioid Receptors Reduce Potassium K Current via Activation of G s Proteins in Cultured Hippocampal Neurons. Journal of Neurophysiology 84: 2356–2364.

74. McLennan GP, Kiss A, Miyatake M, Belcheva MM, Chambers KT, Pozek JJ, et al. (2008): Kappa opioids promote the proliferation of astrocytes via Gbetagamma and beta-arrestin 2-dependent MAPK-mediated pathways. J Neurochem 107: 1753–1765.

75. Potter DN, Damez-Werno D, Carlezon WA, Cohen BM, Chartoff EH (2011): Repeated Exposure to the κ-Opioid Receptor Agonist Salvinorin A Modulates Extracellular Signal-Regulated Kinase and Reward Sensitivity. Biological Psychiatry 70: 744–753.

76. Bruchas MR, Schindler AG, Shankar H, Messinger DI, Miyatake M, Land BB, et al. (2011): Selective p38α MAPK Deletion in Serotonergic Neurons Produces Stress Resilience in Models of Depression and Addiction. Neuron 71: 498–511.

77. Bruchas MR, Macey TA, Lowe JD, Chavkin C (2006): Kappa Opioid Receptor Activation of p38 MAPK Is GRK3- and Arrestin-dependent in Neurons and Astrocytes*. Journal of Biological Chemistry 281: 18081–18089.

78. Yoshizawa K, Narita M, Saeki M, Narita M, Isotani K, Horiuchi H, et al. (2011): Activation of extracellular signal-regulated kinase is critical for the discriminative stimulus effects induced by U-50,488H. Synapse 65: 1052–1061.

79. Li J-G, Luo L-Y, Krupnick JG, Benovic JL, Liu-Chen L-Y (1999): U50,488H-induced Internalization of the Human κ Opioid Receptor Involves a β-Arrestin- and Dynamin-dependent Mechanism: κ RECEPTOR INTERNALIZATION IS NOT REQUIRED FOR MITOGEN-ACTIVATED PROTEIN KINASE ACTIVATION*. Journal of Biological Chemistry 274: 12087–12094.

80. Ehrich JM, Messinger DI, Knakal CR, Kuhar JR, Schattauer SS, Bruchas MR, et al. (2015): Kappa Opioid Receptor-Induced Aversion Requires p38 MAPK Activation in VTA Dopamine Neurons. J Neurosci 35: 12917–12931.

81. Fadok JP, Dickerson TMK, Palmiter RD (2009): Dopamine Is Necessary for Cue-Dependent Fear Conditioning. J Neurosci 29: 11089–11097.

82. Janak PH, Tye KM (2015): From circuits to behaviour in the amygdala. Nature 517: 284–292.

83. Tang W, Kochubey O, Kintscher M, Schneggenburger R (2020): A VTA to Basal Amygdala Dopamine Projection Contributes to Signal Salient Somatosensory Events during Fear Learning. J Neurosci 40: 3969–3980.

84. Hyman SE, Malenka RC, Nestler EJ (2006): NEURAL MECHANISMS OF ADDICTION: The Role of Reward-Related Learning and Memory. Annu Rev Neurosci 29: 565–598.

85. Robinson TE, Berridge KC (2008): The incentive sensitization theory of addiction: some current issues. Phil Trans R Soc B 363: 3137–3146.

86. See RE (2002): Neural substrates of conditioned-cued relapse to drug-seeking behavior. Pharmacology Biochemistry and Behavior 71: 517–529.

87. Milton AL, Everitt BJ (2012): The persistence of maladaptive memory: Addiction, drug memories and anti-relapse treatments. Neuroscience & Biobehavioral Reviews 36: 1119–1139.

88. Lueptow LM, Shashkova EC, Miller MG, Evans CJ, Cahill CM (2020): Insights into the Neurobiology of Craving in Opioid Use Disorder. Curr Anesthesiol Rep 10: 378–387.

