## Supplementary Information for "Divergent roles of orexin and dynorphin in modulating VTA dopamine projections to the BLA after morphine exposure"

#### Supplementary Figures and Figure legends

Figure S1

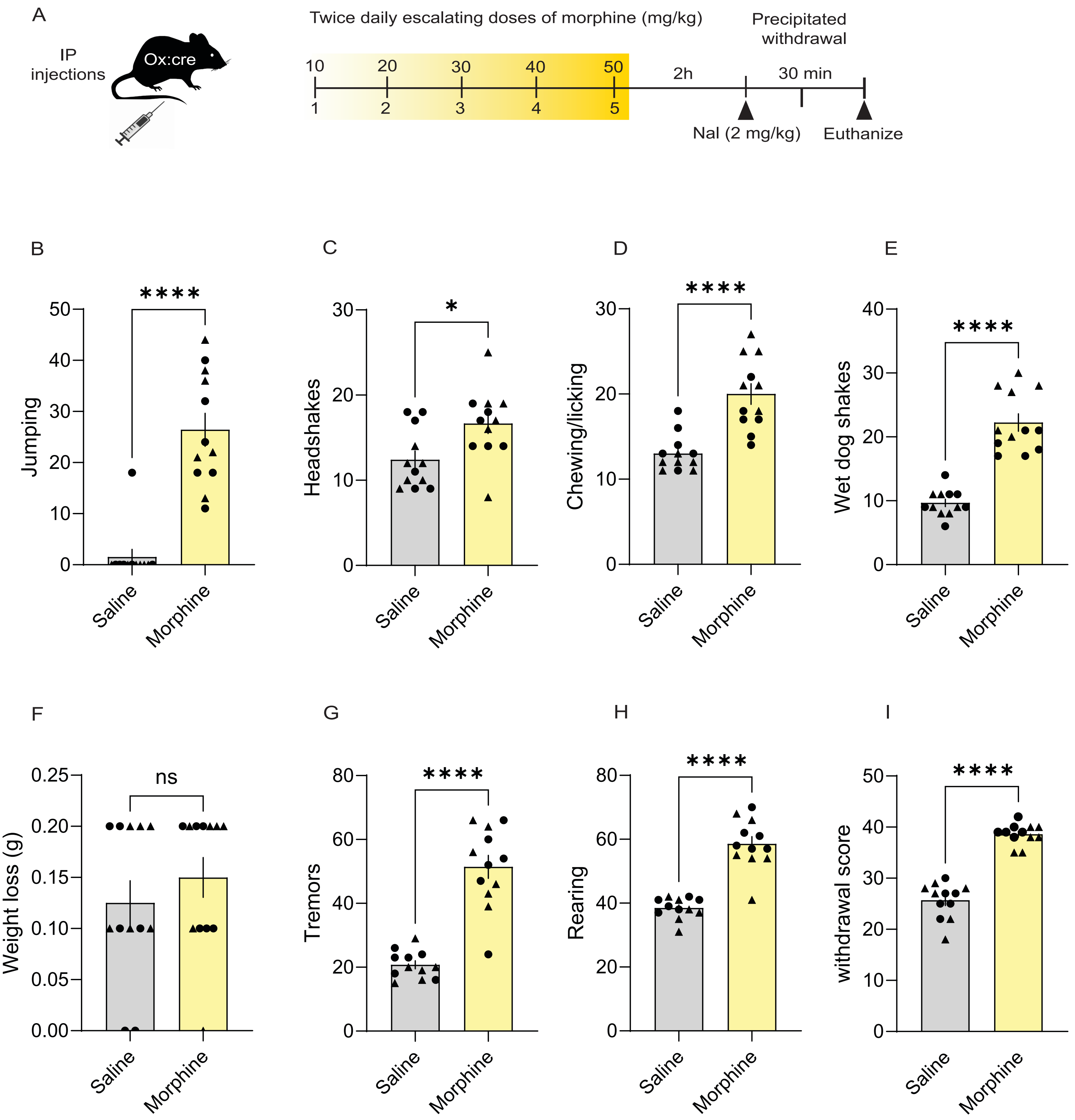

A
IP injections
Ox:cre
Twice daily escalating doses of morphine (mg/kg)
10 20 30 40 50
1 2 3 4 5
2h
Precipitated withdrawal
30 min
Nal (2 mg/kg)
Euthanize
B
Jumping
****
Saline
Morphine
C
Headshakes
*
Saline
Morphine
D
Chewing/licking
****
Saline
Morphine
E
Wet dog shakes
****
Saline
Morphine
F
Weight loss (g)
ns
Saline
Morphine
G
Tremors
****
Saline
Morphine
H
Rearing
****
Saline
Morphine
I
withdrawal score
****
Saline
Morphine

Figure S2

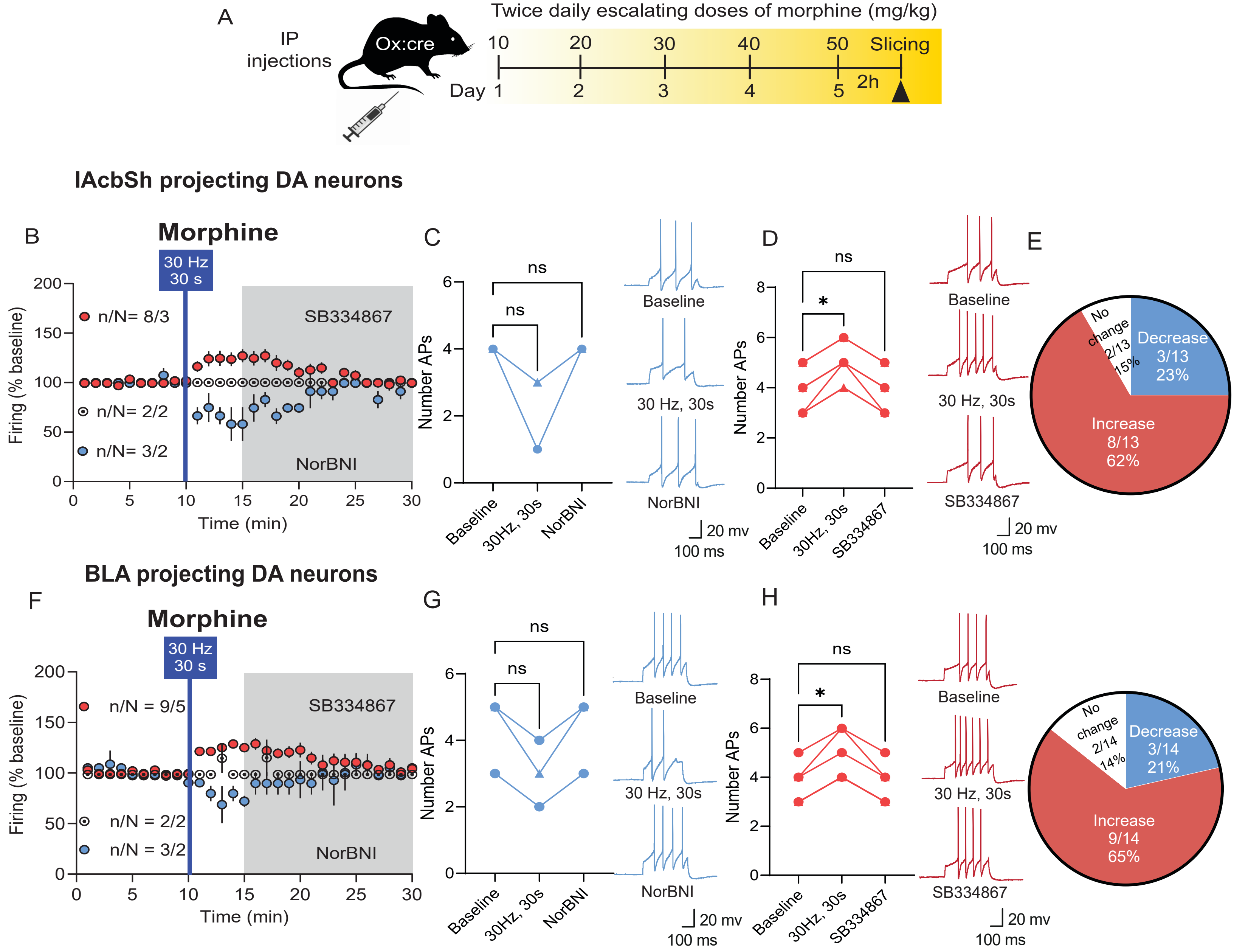

A
IP injections
Ox:cre
Twice daily escalating doses of morphine (mg/kg)
10
20
30
40
50
Slicing
Day 1
2
3
4
5
2h
lAcbSh projecting DA neurons
B
Morphine
30 Hz 30 s
SB334867
NorBNI
n/N= 8/3
n/N= 2/2
n/N= 3/2
Firing (% baseline)
Time (min)
C
ns
ns
Number APs
Baseline
30Hz, 30s
NorBNI
Baseline
30 Hz, 30s
NorBNI
20 mv
100 ms
D
ns
*
Number APs
Baseline
30Hz, 30s
SB334867
Baseline
30 Hz, 30s
SB334867
20 mv
100 ms
E
No change 2/13 15%
Decrease 3/13 23%
Increase 8/13 62%
BLA projecting DA neurons
F
Morphine
30 Hz 30 s
SB334867
NorBNI
n/N = 9/5
n/N = 2/2
n/N = 3/2
Firing (% baseline)
Time (min)
G
ns
ns
Number APs
Baseline
30Hz, 30s
NorBNI
Baseline
30 Hz, 30s
NorBNI
20 mv
100 ms
H
ns
*
Number APs
Baseline
30Hz, 30s
SB334867
Baseline
30 Hz, 30s
SB334867
20 mv
100 ms
No change 2/14 14%
Decrease 3/14 21%
Increase 9/14 65%

Figure S3

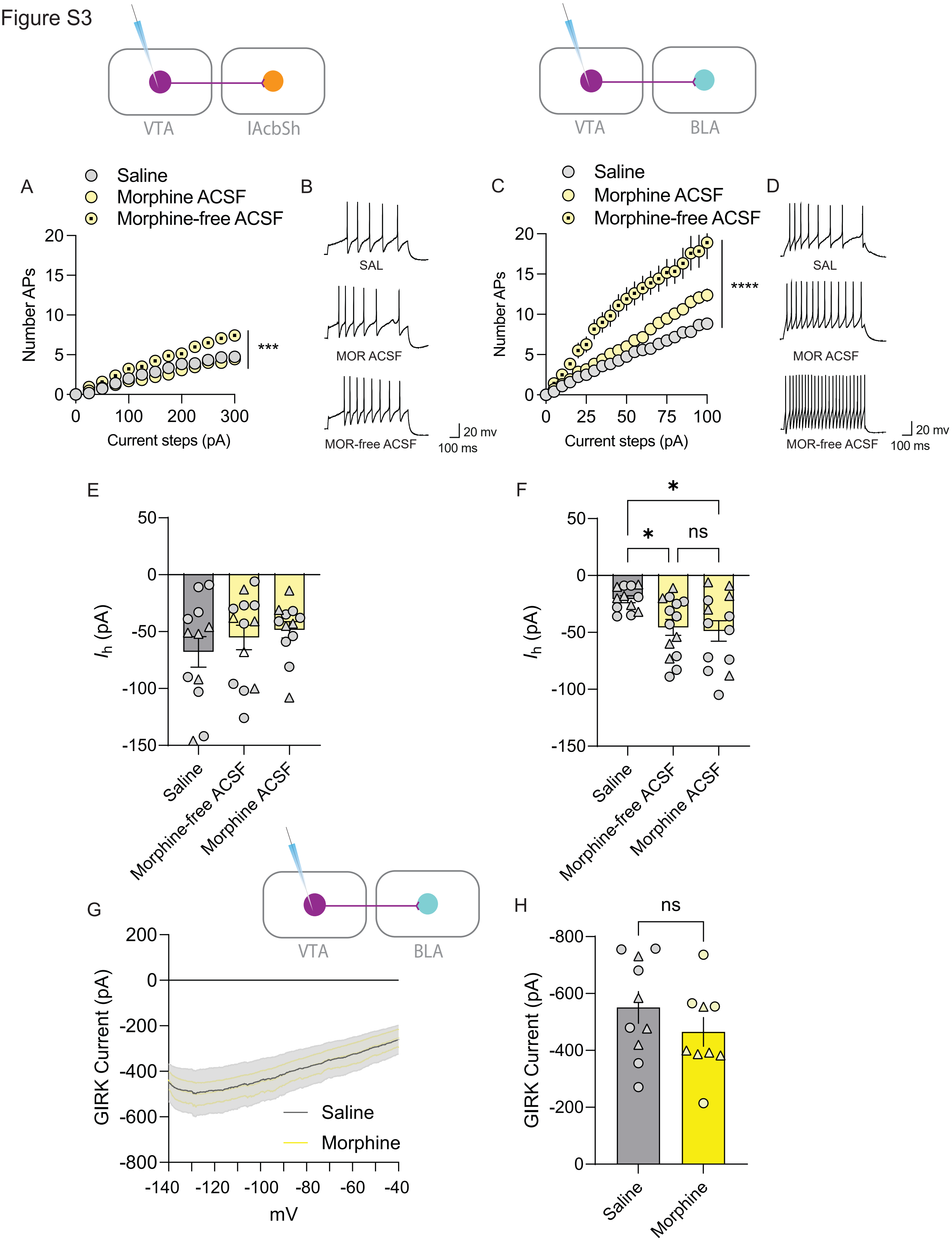
VTA
lAcbSh
VTA
BLA
A
Saline
Morphine ACSF
Morphine-free ACSF
Number APs
Current steps (pA)
***
B
SAL
MOR ACSF
MOR-free ACSF
20 mv
100 ms
C
Saline
Morphine ACSF
Morphine-free ACSF
Number APs
Current steps (pA)
****
D
SAL
MOR ACSF
MOR-free ACSF
20 mv
100 ms
E
$I_h$ (pA)
Saline
Morphine-free ACSF
Morphine ACSF
F
*
*
ns
$I_h$ (pA)
Saline
Morphine-free ACSF
Morphine ACSF
G
VTA
BLA
GIRK Current (pA)
Saline
Morphine
mV
H
ns
GIRK Current (pA)
Saline
Morphine

Figure S4

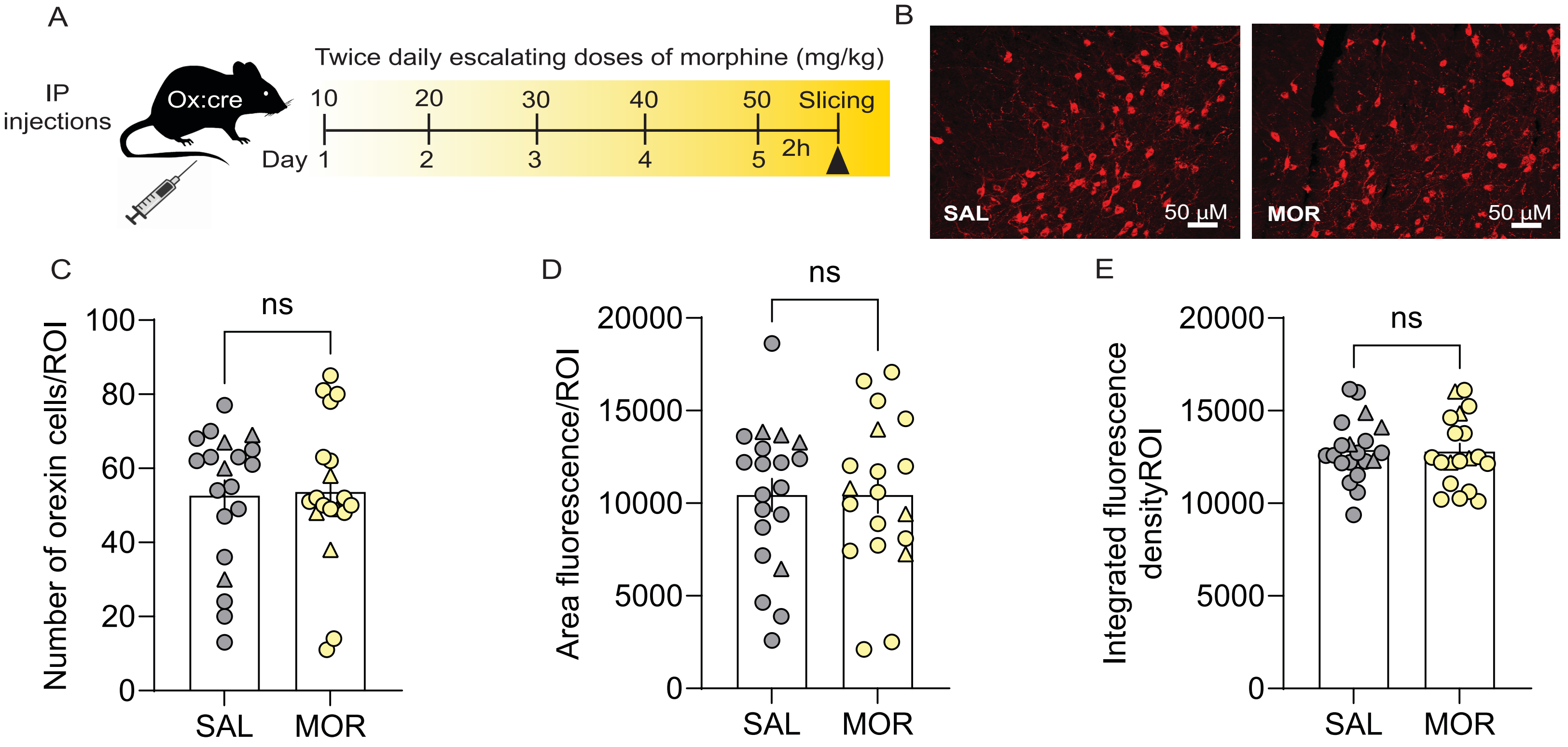
A
IP injections
Ox:cre
Twice daily escalating doses of morphine (mg/kg)
10
20
30
40
50
Slicing
Day 1
2
3
4
5
2h
B
SAL
50 µM
MOR
50 µM
C
ns
Number of orexin cells/ROI
100
80
60
40
20
0
SAL
MOR
D
ns
Area fluorescence/ROI
20000
15000
10000
5000
0
SAL
MOR
E
ns
Integrated fluorescence densityROI
20000
15000
10000
5000
0
SAL
MOR

Figure S5

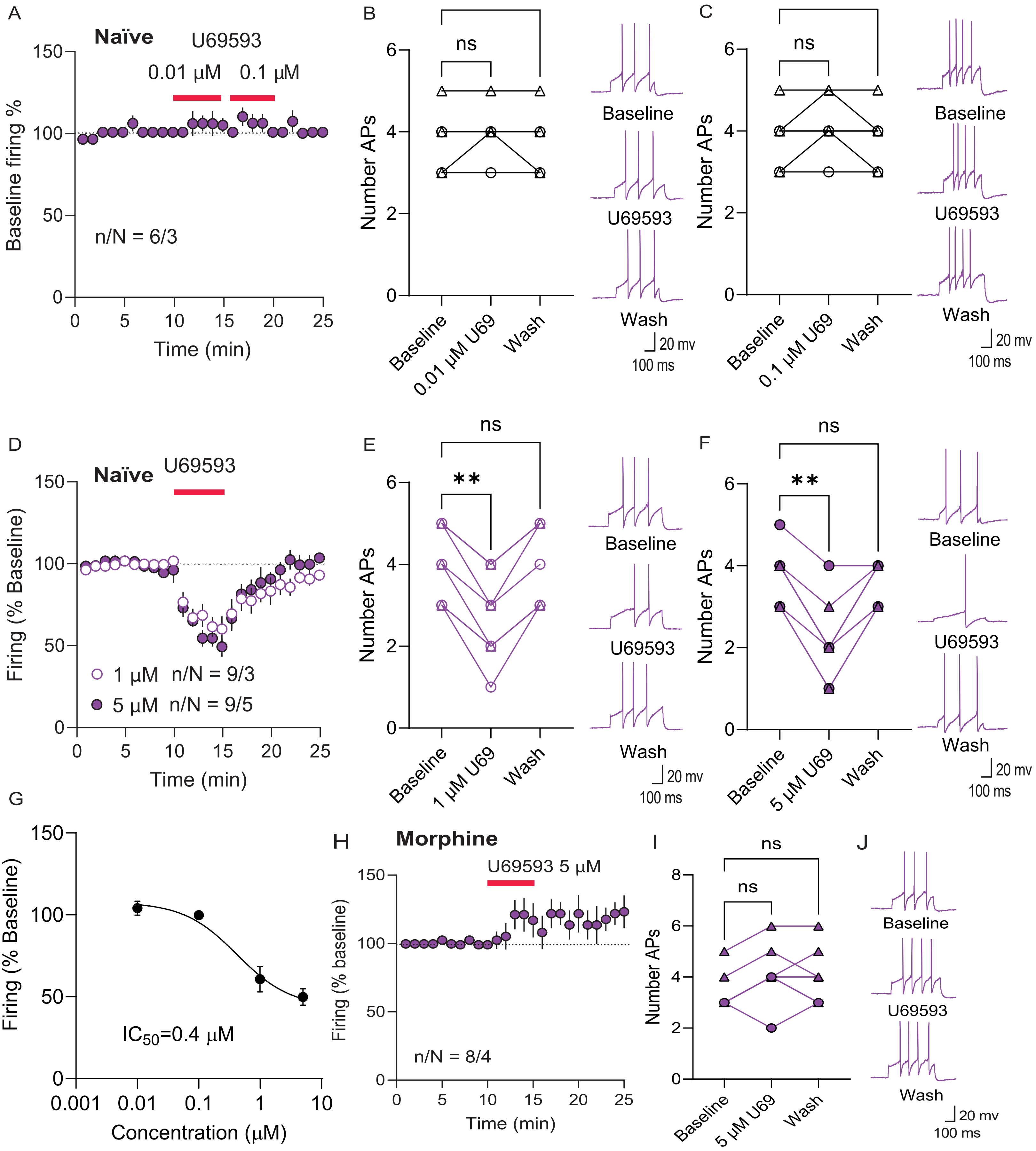

A
Naïve
U69593
0.01 μM
0.1 μM
Baseline firing %
n/N = 6/3
Time (min)
B
ns
ns
Number APs
Baseline
0.01 μM U69
Wash
Baseline
U69593
Wash
20 mv
100 ms
C
ns
ns
Number APs
Baseline
0.1 μM U69
Wash
Baseline
U69593
Wash
20 mv
100 ms
D
Naïve
U69593
Firing (% Baseline)
1 μM n/N = 9/3
5 μM n/N = 9/5
Time (min)
E
ns
**
Number APs
Baseline
1 μM U69
Wash
Baseline
U69593
Wash
20 mv
100 ms
F
ns
**
Number APs
Baseline
5 μM U69
Wash
Baseline
U69593
Wash
20 mv
100 ms
G
Firing (% Baseline)
IC50=0.4 μM
Concentration (μM)
H
Morphine
U69593 5 μM
Firing (% baseline)
n/N = 8/4
Time (min)
I
ns
ns
Number APs
Baseline
5 μM U69
Wash
J
Baseline
U69593
Wash
20 mv
100 ms

Figure S6

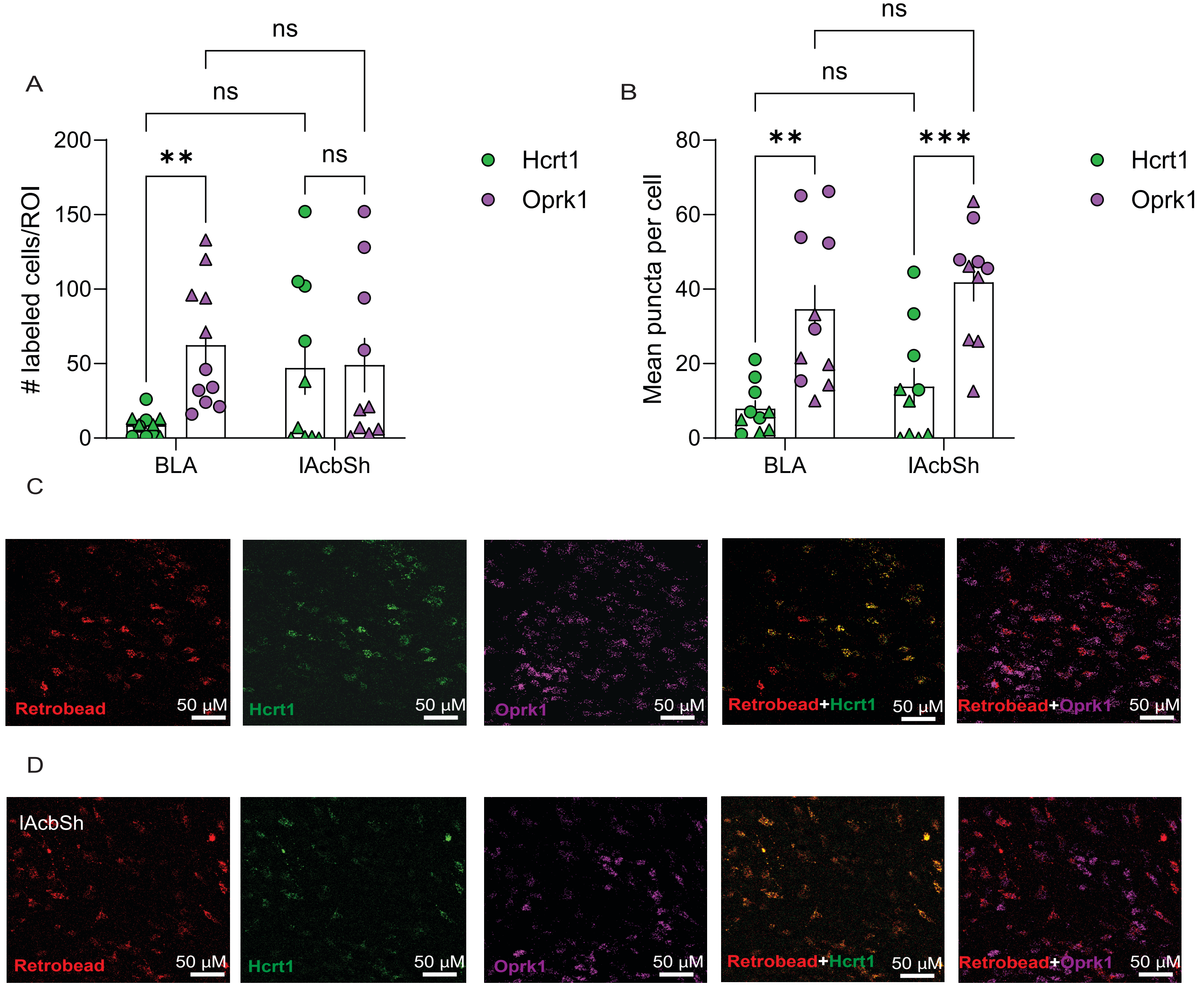

A
ns
ns
ns
**
200
150
100
50
0
### labeled cells/ROI
BLA
lAcbSh
Hcrt1
Oprk1
B
ns
ns
**
***
80
60
40
20
0
Mean puncta per cell
BLA
lAcbSh
Hcrt1
Oprk1
C
Retrobead
50 μM
Hcrt1
50 μM
Oprk1
50 μM
Retrobead+Hcrt1
50 μM
Retrobead+Oprk1
50 μM
D
lAcbSh
Retrobead
50 μM
Hcrt1
50 μM
Oprk1
50 μM
Retrobead+Hcrt1
50 μM
Retrobead+Oprk1
50 μM

Figure S7

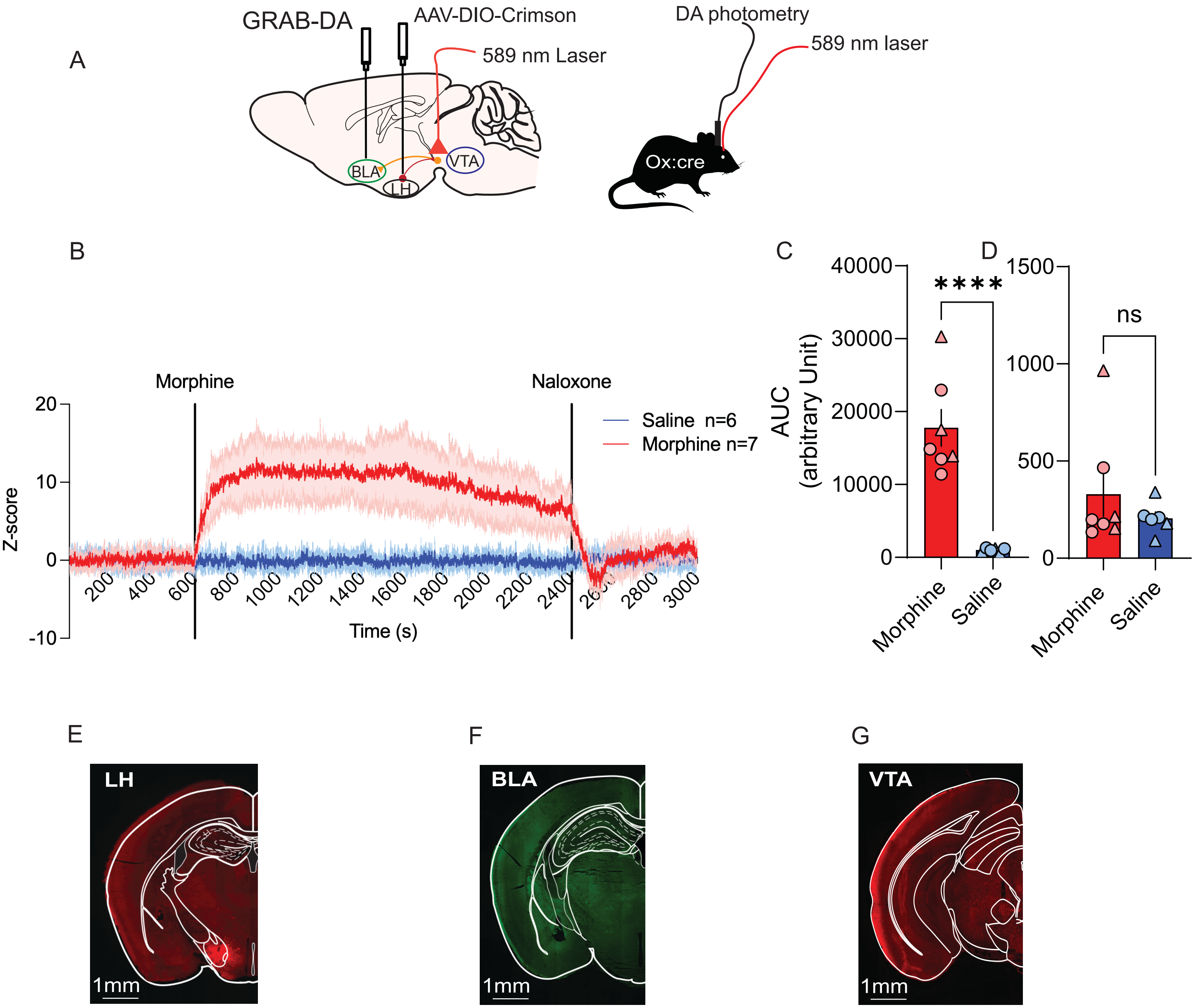
A
GRAB-DA
AAV-DIO-Crimson
589 nm Laser
BLA
LH
VTA
DA photometry
589 nm laser
Ox:cre
B
Morphine
Naloxone
Saline n=6
Morphine n=7
Z-score
Time (s)
C
D
AUC
(arbitrary Unit)
****
ns
Morphine
Saline
E
LH
1mm
F
BLA
1mm
G
VTA
1mm

Figure S8

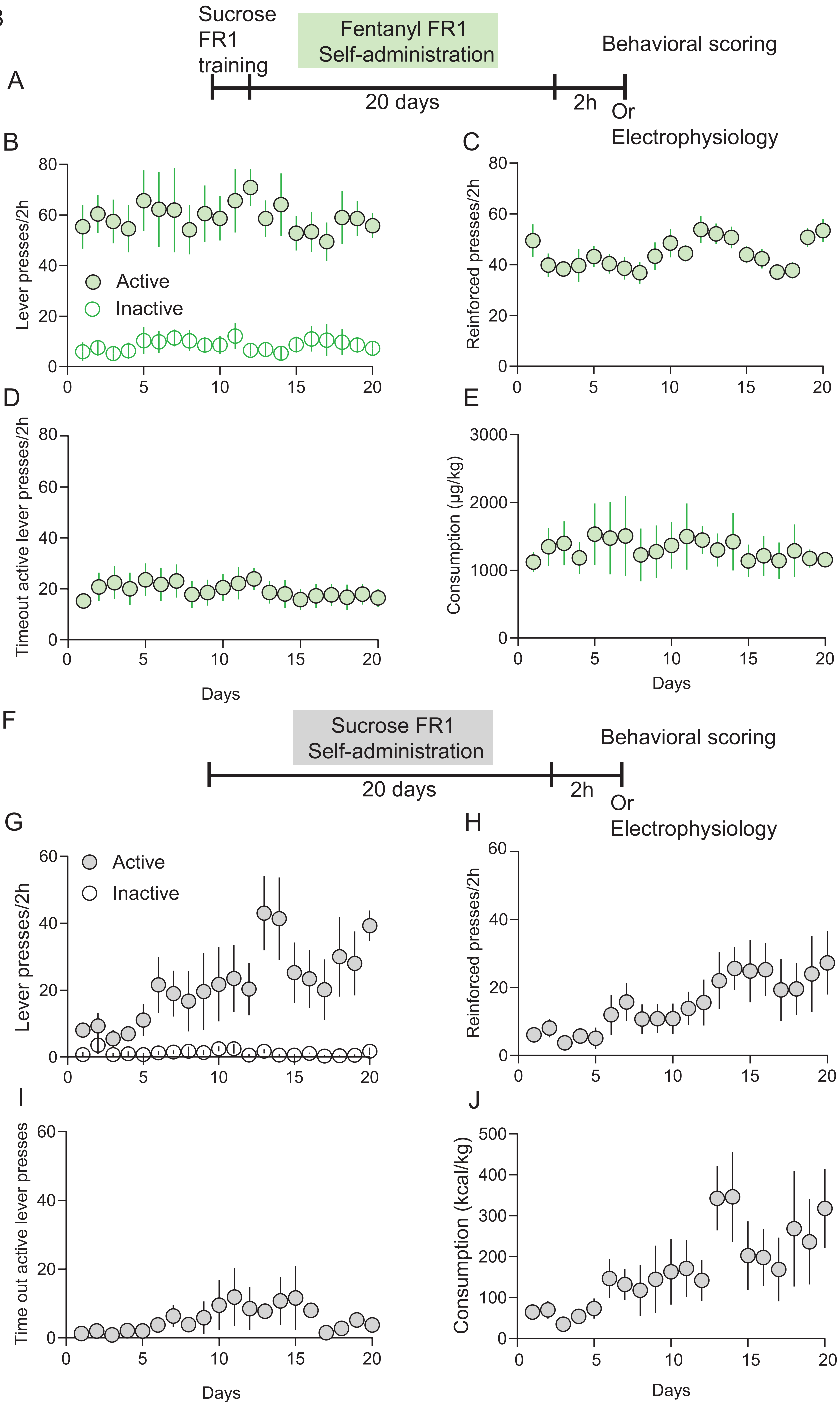
A
Sucrose FR1 training
Fentanyl FR1 Self-administration
Behavioral scoring
20 days
2h
Or Electrophysiology
B
Lever presses/2h
Active
Inactive
C
Reinforced presses/2h
D
Timeout active lever presses/2h
Days
E
Consumption (µg/kg)
Days
F
Sucrose FR1 Self-administration
Behavioral scoring
20 days
2h
Or Electrophysiology
G
Lever presses/2h
Active
Inactive
H
Reinforced presses/2h
I
Time out active lever presses
Days
J
Consumption (kcal/kg)
Days

Figure S9

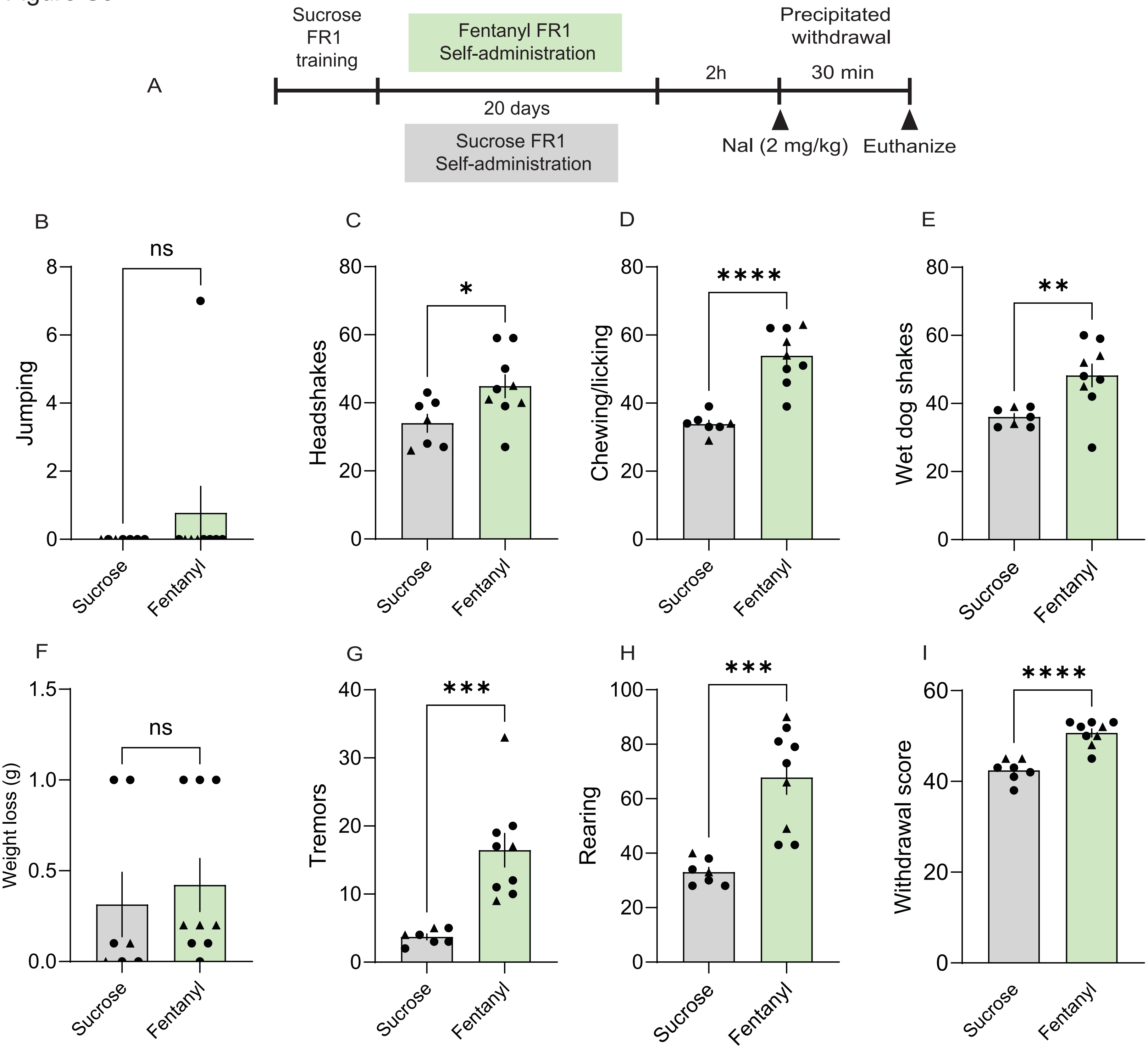
A
Sucrose
FR1
training
Fentanyl FR1
Self-administration
20 days
Sucrose FR1
Self-administration
2h
Precipitated
withdrawal
30 min
Nal (2 mg/kg)
Euthanize
B
ns
Jumping
8
6
4
2
0
Sucrose
Fentanyl
C
*
Headshakes
80
60
40
20
0
Sucrose
Fentanyl
D
****
Chewing/licking
80
60
40
20
0
Sucrose
Fentanyl
E
**
Wet dog shakes
80
60
40
20
0
Sucrose
Fentanyl
F
ns
Weight loss (g)
1.5
1.0
0.5
0.0
Sucrose
Fentanyl
G
***
Tremors
40
30
20
10
0
Sucrose
Fentanyl
H
***
Rearing
100
80
60
40
20
0
Sucrose
Fentanyl
I
****
Withdrawal score
60
40
20
0
Sucrose
Fentanyl

**Supplementary Figure 1. Naloxone-precipitated withdrawal occurs with escalating repeated morphine treatment (supporting figure 1).**

(A) Schematic of twice-daily escalating morphine injections and timecourse of precipitated withdrawal measurement. Naloxone led to significant increases in (B) jumping ($t_{(22)}$ = 7.022, p<0.0001), (C) head shakes ($t_{(22)}$ = 2.73, p=0.012), (D) chewing and licking behaviours ($t_{(22)}$ = 5.170, p<0.0001), (E) wet dog shakes ($t_{(22)}$ = 8.51, p<0.0001), (G) tremors ($t_{(22)}$ = 8.048, p<0.0001), and (H) rearing ($t_{(22)}$ = 8.27, p<0.0001), but no change in (F) weight ($t_{(22)}$ =0.856, p = 0.40) in morphine-dependent compared to saline mice. (I) Cumulative withdrawal scores in orexin$^{cre}$ mice indicated naloxone precipitated withdrawal was more severe in mice administered morphine compared to saline ($t_{(22)}$ = 7.79, p<0.0001). 12 mice per saline or morphine group. Data presented as mean ± SEM. ∗p < 0.05, ∗∗p < 0.01, ∗∗∗p < 0.001, ∗∗∗p < 0.0001.

**Supplemental Figure 2. Repeated morphine increases firing of BLA-projecting VTA$^{DA}$ neurons in response to optical stimulation of LH$_{ox/dyn}$ inputs, but does not change the neuromodulatory capacity of LH$_{ox/dyn}$ on lAcbSh-projecting VTA$^{DA}$ neurons (Supporting figure 2).**

(A) Timeline of escalating twice-daily morphine treatment.

(B) Time course of evoked firing before and after photoactivation of LH$_{ox/dyn}$ inputs onto lAcbSh-projecting VTA$^{DA}$ neurons of morphine-treated mice (n/N=13/7).

(C, D) Number of action potentials and sample traces that showed decreased (J; n/N=3/2, $F_{(2, 2)}$=4, p=0.27) or increased (K; n/N=8/3, $F_{(2, 12)}$= 64, p=0.010) responses to $LH_{ox/dyn}$ optical stimulation in morphine-treated mice.

(E) In morphine-treated mice, while photoactivation of $LH_{ox/dyn}$ inputs primarily increases firing rate of lAcbSh-projecting $VTA^{DA}$ neurons, it is not different from saline-treated mice in Figure 2G ($\chi^2$=0.21, p=0.65).

(F) Time course of evoked firing before and after photoactivation of $LH_{ox/dyn}$ inputs in BLA-projecting $VTA^{DA}$ neurons of morphine-treated mice (n/N=14/9).

(G, H) Number of action potentials and sample traces that showed decreased (J; n/N=3/2, $F_{(2, 4)}$= 16, p=0.13) or increased (K; n/N=9/5, $F_{(2, 12)}$= 64, p=0.010) responses to $LH_{ox/dyn}$ optical stimulation in morphine-treated mice.

(I) In morphine-treated mice, photoactivation of $LH_{ox/dyn}$ inputs primarily increased firing rate of BLA-projecting $VTA^{DA}$ neurons compared to saline-treated mice in Figure 2R ($\chi^2$=5.24, p=0.022).

Data presented as mean ± SEM. $p < 0.05$, $p < 0.05$.

**Supplementary Figure 3. Morphine treatment increases excitability of $VTA^{DA}$ neurons that project to either the BLA or lAcbSh (Supporting figure 2).**

(A) Frequency-current (F-I) plot of action potentials in lAcbSh-projecting $VTA^{DA}$ neurons from saline or morphine treated mice in the presence or absence of continued morphine. Drug x current step interaction: $F_{(5.52, 99.5)}$ = 6.85, P<0.0001, main effect drug: $F_{(2, 36)}$ = 11.65, P=0.0001, main effect current step: $F_{(2.76,\ 99.5)} = 277.8$, $P<0.0001$. Dunnet’s multiple comparisons test indicated significant differences between saline (n/N=12/8) and morphine-free ACSF (n/N=13/7) at all current steps except 25 pA ($p<0.05$) but no differences between saline and morphine ACSF(n/N=12/7).

(B) Example traces from the 300-pA step in lAcbSh-projecting $VTA^{DA}$ neurons.

(C) F-I plot of action potentials in BLA-projecting $VTA^{DA}$ neurons from saline or morphine treated mice in the presence or absence of continued morphine. Drug x current step interaction: $F_{(4.66,\ 79.18)} = 9.58$, $P<0.0001$, main effect drug: $F_{(2,\ 34)} = 18.60$, $P<0.0001$, main effect current step: $F_{(2.33,\ 79.18)} = 201.4$, $P<0.0001$. Dunnet’s multiple comparisons test indicated significant differences between saline (n/N=12/8) and morphine-free ACSF (n/N=14/9) at all current steps except 5 and 10 pA ($p<0.05$) and only differences in saline and morphine ACSF (n/N=13/10) at 65-100 pA ($p < 0.05$).

(D) Example traces from the 100-pA step in BLA-projecting $VTA^{DA}$ neurons.

(E) $I_h$ recorded from lAcbSh-projecting $VTA^{DA}$ neurons from saline mice was not significantly different from that of morphine treated mice in the presence or absence of continued morphine. One way ANOVA: $F_{(2,34)} = 0.82$, $P = 0.45$.

(F) $I_h$ recorded from BLA-projecting $VTA^{DA}$ neurons from saline mice was significantly different from that of morphine treated mice in the presence ($p = 0.016$) or absence ($p = 0.31$) of continued morphine. One way ANOVA: $F_{(2,36)} = 5.89$, $P = 0.017$, Sidak’s post hoc test.

(G) GIRK current from BLA-projecting $VTA^{DA}$ neurons from saline- and morphine-treated mice (n/N=9/5).

(H) Maximum GIRK current of BLA-projecting $VTA^{DA}$ neurons is not different between saline-treated compared to morphine-treated mice ($t_{(17)}$= 1.13, P=0.27).

Data presented as mean ± SEM. ∗p < 0.05.

**Supplementary Figure 4. Repeated morphine did not increase orexin expression (Supporting Figure 3).**

(A) Timeline of escalating twice-daily morphine treatment.

(B) Representative immunofluorescence images of orexin neurons in the LH from mice treated with saline (left) or morphine (right) twice daily for 5 days.

(C) Number of orexin-positive cells per region of interest (ROI) is not different between saline- and morphine-treated mice ($t_{(37)}$=0.16, p=0.87).

(D) Mean fluorescence area per ROI is not different between saline- and morphine-treated mice ($t_{(37)}$= 0.0004, p=0.99).

(E) Integrated fluorescence density per ROI is not different between saline- and morphine-treated mice ($t_{(37)}$= 0.14, p=0.89).

Data presented as mean ± SEM. ∗p < 0.05, ∗∗∗p < 0.001. Scale bars, 50 μm.

**Supplementary Figure 5. The KOR agonist U69593 inhibited the firing rate of BLA-projecting $VTA^{DA}$ neurons in naïve mice (Supporting Figure 3).**

(A) Time course of evoked firing before and after bath application of U69593 (0.01 and 0.1 μM) in BLA-projecting $VTA^{DA}$ neurons from naïve mice (n/N=6/3).

(B) Number of action potentials was not different after U69593 (0.01 μM) bath application ($F_{(2,10)}$ =1, p=0.40) to BLA-projecting $VTA^{DA}$ neurons. Sample traces of evoked action potentials before, during, and after U69593 (0.01 μM).

(C) Number of action potentials was not different after U69593 (0.1 μM) bath application ($F_{(2,10)}$ = 2.50, p=0.13) applied to BLA-projecting $VTA^{DA}$ neurons. Sample traces of evoked action potentials before, during, and after U69593 (0.1 μM).

(D) Time course of evoked firing before and after bath application of U69593 (1 (open circles) and 5 μM (filled circles)) to BLA-projecting $VTA^{DA}$ neurons from naïve mice (n/N=9/5).

(E) Number of action potentials decreased with bath application of U69593 (1 μM) to BLA-projecting $VTA^{DA}$ neurons ($F_{(2,16)}$ = 53.20, p=0.0019). Sample traces of evoked action potentials before, during, and after U69593 (1 μM).

(F) Number of action potentials decreased after U69593 (5 μM) bath application to BLA-projecting $VTA^{DA}$ neurons ($F_{(2,16)}$ = 45.49, p=0019). Sample traces of evoked action potentials before, during, and after U69593 (5 μM).

(G) Dose–response curve of the KOR agonist U69593 on evoked firing fit with a non-linear inhibitor vs response curve indicating an $IC_{50}$ of 0.42 μM.

(H) Time course of evoked firing before and after bath application of U69593 in BLA-projecting $VTA^{DA}$ neurons from morphine-treated mice (n/N=8/4).

(I) In morphine-treated mice, U69593 did not alter the number of action potentials of BLA-projecting VTA$^{DA}$ neurons (n/N=8/4) ($F_{(2, 14)}$= 4.43, p=0.067).

(J) Sample traces of evoked action potentials.

Data presented as mean ± SEM. $**p < 0.01$.

**Supplementary Figure 6. BLA-projecting VTA$^{DA}$ neurons exhibit higher *Oprk1* than *Hcrt1* mRNA expression (Supporting Figure 3).**

(A) The number of BLA-projecting VTA$^{DA}$ neurons expressing *Oprk1* mRNA was significantly greater than those with *Hcrt1* expression.Two-way ANOVA (region × marker interaction: $F_{(1,38)}$=0.598, P=0.089; main effect of markers: $F_{(1,38)}$=4.11, P=0.049; main effect of region: $F_{(1,38)}$=0.59, P=0.45). Fisher's posthoc test revealed that there was greater expression of Oprk1 mRNA in BLA-projecting VTA$^{DA}$ neurons (p=0.007), whereas there was no significant difference in expression of either receptor in lAcbSh-projecting VTA$^{DA}$ neurons (p=0.92).

(B) Averaged Oprk1 puncta per cell were significantly higher than Hcrt1 in VTA$^{DA}$ neurons projecting to both the BLA and lAcbSh. Two-way ANOVA (region × marker interaction: $F_{(1,22)}$ = 0.056, P=0.86; main effect of markers: $F_{(1,22)}$=50.96, P<0.0001; main effect of region: $F_{(1,20)}$ = 2.02, P=0.171). Tukey's posthoc test revealed that averaged Oprk1 puncta per cell was greater than Hcrt1 in VTA$^{DA}$ neurons projecting to both the BLA (p=0.0054) and lAcbSh (p=0.0004).

(C) 20x confocal images of VTA neurons projecting to the BLA of orexin$^{cre}$ mice showing *Hcrt1* (green), retrobead (red), *Oprk1* (purple), and merged co-expression. Scale bar =50 μm

(D) 20x confocal images of VTA neurons projecting to the lAcbSh of $orexin^{cre}$ mice showing *Hcrt1* (green), retrobead (red), *Oprk1* (purple), and merged co-expression. Scale bar =50 μm

Data presented as mean ± SEM. ∗∗p < 0.01.

**Supplementary Figure 7. Morphine increases $BLA^{DA}$ in morphine dependent mice (Supporting Figure 5).**

(A) Schematic of viral strategy whereby $GRAB_{DA}$ and a photometry ferrule are expressed in the BLA, chrimson is expressed in $LH_{ox/dyn}$ neurons, and the optic ferrule is implanted to the VTA.

(B) Time course of Z-scored photometry responses showing that systemic morphine (50 mg/kg) increased BLA dopamine, which was suppressed by naloxone (2 μM) treatment in morphine-dependent mice.

(C) Morphine (N = 7) evokes greater BLA dopamine that saline (N = 6) in morphine-dependent mice, measured by area under the curve (AUC) (unpaired two-tailed t-test: $t_{(11)} = 6.13$, $P < 0.0001$).

(D) Naloxone-induced suppression of BLA dopamine in morphine-dependent mice administered morphine was not significantly different from mice administered saline (unpaired two-tailed t-test: $t_{(11)} = 0.97$, $P = 0.35$).

(E) Example image of the LH chrimson injection site.

(F) Example image of $GRAB_{DA}$ expression and photometry ferule placement in the BLA.

(G) Example image of chrimson expression in a slice containing the VTA.

Data presented as mean ± SEM. ∗∗∗p < 0.0001.

**Supplementary Figure 8. Oral fentanyl and sucrose self-administration (Supporting Figure 6)**.

(A) Experimental timeline of fentanyl self-administration under an FR1 schedule.

(B) Active (filled circles) and inactive (open circles) lever presses across 20 fentanyl self-administration sessions. Initial sucrose fade is not shown. During FR1 training, mice made more presses on the active lever than the inactive lever indicating discrimination of fentanyl administration (main effect of lever: $F_{(1,20)}$=41.48, p<0.0001). (N = 5 male and 6 female mice).

(C) Number of reinforced presses across sessions changed over the time course (RM one way ANOVA: $F_{(1, 20)}$ =41.48, p=0.0001).

**(**D) Time out lever presses during self-administration did not change over the time course (RM one way ANOVA: $F_{(19, 190)}$ =0.87, p=0.62).

(E) Total fentanyl consumption per day did not change over the time course (RM one way ANOVA: $F_{(19, 190)}$ =0.50, p=0.96).

(F) Experimental timeline of sucrose (10%) self-administration.

(G) Active and inactive lever presses across sucrose self-administration sessions. During FR1 training, mice made more presses on the active lever delivering sucrose than the inactive (main effect of lever: $F_{(1,14)=}$11.61, p=0.0043) (N = 4 male and 3 female mice).

(H) Number of reinforced presses across FR1 sessions changed over the time course (RM one way ANOVA: $F_{(7, 120)}$ =17.43, p=0.0028).

(I) Time out lever presses during sucrose self-administration.

(J) Total sucrose consumption per day changed over the time course (RM one way ANOVA: $F_{(7, 120)}$ =8.91, p=0.0055).

Data presented as mean ± SEM. ∗p < 0.05, ∗∗p < 0.01, ∗∗∗p < 0.001, ∗∗∗p < 0.0001.

**Supplementary Figure 9. Naloxone-precipitated withdrawal signs following fentanyl self-administration (Supporting Figure 6).**

(A) Two hours after the last self-administration session, naloxone was injected to precipitated withdrawal. Naloxone led to significant increases in (C) head shakes ($t_{(14)}$ = 2.43, p=0.029), (D) chewing and licking behaviors ($t_{(14)}$=6.11, p<0.0001), (E) wet-dog shakes ($t_{(14)}$ =3.12, p=0.0076), (G) tremors ($t_{(14)}$ = 4.47, p=0.0005), and (H) rearing ($t_{(14)}$ = 4.82, p=0.0003), in mice that underwent fentanyl compared to sucrose self-administration. No significant differences were observed in (B) jumping ($t_{(14)}$ = 0.87, p=0.99), (F) weight loss ($t_{(14)}$ = 0.47, p=0.36). (M) Cumulative withdrawal scores demonstrate that naloxone-precipitated withdrawal was more severe in fentanyl (N=9) compared to sucrose self-administering mice (N=7; $t_{(14)}$ = 6.25, p<0.0001).

Data presented as mean ± SEM. ∗p < 0.05, ∗∗p < 0.01, ∗∗∗p < 0.001, ∗∗∗p < 0.0001.

**Supplemental Table 1.** Composite scoring for withdrawal behaviour in adult morphine or fentanyl exposed mice.

| Behaviour | Number[a] | Score |
|---|---|---|
| **Jumping** | 1-5 | 1 |
| | 6-10 | 2 |
| | >10 | 3 |
| **Headshakes** | 1-2 | 1 |
| | 3-4 | 2 |
| | >4 | 3 |
| **Wet-dog shakes** | 1-2 | 1 |
| | 3-4 | 2 |
| | >4 | 3 |
| **Chewing/licking** | 1-9 | 1 |
| | 10-15 | 2 |
| | >15 | 3 |
| **Weight loss** | 0.1 | 1 |
| | 0.2-0.4 | 2 |
| | >0.4 | 3 |
| **Tremors** | Absent | 0 |
| | Present | 1 |

[a]Number of occurrences and absence/presence per 5 min interval. Total score is calculated by taking the sum of all weighted scores over the whole 30 min.
